# Early and Late Effects of Maternal Experience on Hippocampal Neurogenesis, Microglia, and the Circulating Cytokine Milieu

**DOI:** 10.1101/388009

**Authors:** Rand S Eid, Jessica A Chaiton, Stephanie E Lieblich, Tamara S Bodnar, Joanne Weinberg, Liisa AM Galea

## Abstract

The maternal brain displays considerable plasticity, and motherhood is associated with changes in affective and cognitive function. Motherhood can alter the trajectory of brain ageing, including modifications to neuroplasticity and cognition. Here, we investigated the short- and long-term effects of motherhood on hippocampal neurogenesis, microglial density and morphology, and circulating cytokines, domains known to be altered with age and implicated in cognition and mood. Female rats were bred then euthanized during gestation or at various postpartum timepoints, culminating in middle age, and nulliparous rats served as age-matched controls. Hippocampal neurogenesis was significantly suppressed during gestation and the postpartum period. Interestingly, neurogenesis declined significantly in middle-aged nulliparous rats, but increased in primiparous rats across the same period. Transient postpartum adaptations to the neuroimmune environment of the hippocampus were evidenced, as Iba-1-immunoreactive microglia assumed a de-ramified morphology followed by increased density. Intriguingly, ageing-related changes in circulating cytokines were dependent on parity. These adaptations in neurogenic and immune processes may have ramifications for maternal mood and cognition across the peripartum period and beyond.

## 1. Introduction

Dramatic physiological adaptations occur during pregnancy and the postpartum period to ensure offspring development and survival (Dulac et al., 2014; Hall et al., 2011; Lain and Catalano, 2007; PrabhuDas et al., 2015; Rossant and Cross, 2001). The maternal brain exhibits substantial plasticity, including large-scale volumetric changes (Galea et al., 2000; Hoekzema et al., 2016; Oatridge et al., 2002), alterations in cellular architecture (Leuner and Gould, 2010; Pawluski and Galea, 2006), and hippocampal neurogenesis (Darnaudéry et al., 2007; Leuner et al., 2007; Pawluski and Galea, 2007). While this capacity for plasticity is likely essential for the onset of a repertoire of maternal behaviours (Bridges, 2015), motherhood is also associated with changes in affective function (Bennett et al., 2004; Darcy et al., 2011; O’Hara, 2009), hypothalamic-pituitary-adrenal axis (HPA) regulation (Slattery and Neumann, 2008), and cognition (Cuttler et al., 2011; De Groot et al., 2006; Galea et al., 2000; Kinsley et al., 1999; Pawluski et al., 2006). Interestingly, motherhood may improve the ageing trajectory in terms of cognition (Colucci et al., 2006; Cui et al., 2014; Gatewood et al., 2005), neuroplasticity (Barha et al., 2015; Barha and Galea, 2011; Galea et al., 2018), and cellular aging (Barha et al., 2016), suggesting that the effects of the motherhood on the brain may be long lasting. However, the mechanisms underlying alterations in the ageing maternal brain are not known, but may include modifications in neurogenic or immune processes, both of which were examined in the current study.

The hippocampus produces new neurons across the lifespan (Altman and Das, 1965; Boldrini et al., 2018; Eriksson et al., 1998) and these neurons play a role in certain aspects of learning and memory (Yau et al., 2015), mood regulation (Sahay and Hen, 2007), and the stress response (Snyder et al., 2011). Importantly, several studies found postpartum reductions in hippocampal cell proliferation (Darnaudéry et al., 2007; Leuner et al., 2007; Pawluski and Galea, 2007), cell survival (Pawluski and Galea, 2007) and the density of immature neurons (Workman et al., 2015). Interestingly, motherhood may have contrasting effects on hippocampal neurogenesis with age, as studies have found increased neurogenesis in middle-aged primiparous and multiparous rats relative to age-matched nulliparous controls (Barha et al., 2015; Galea et al., 2018). This finding signifies that parity can have delayed pro-neurogenic effects, thus the current study aimed to determine the timeline by which these changes may emerge.

Adaptations to the maternal immune system are well documented, and necessary for the establishment and maintenance of pregnancy (Aghaeepour et al., 2017; Mor and Cardenas, 2010). In contrast, and despite the growing recognition of plasticity in the maternal brain, there is a paucity of research on potential neuroimmune adaptations with maternal experience. Few studies to date have examined microglia, the innate immune cells of the brain, in pregnant and postpartum rats (Haim et al., 2017; Posillico and Schwarz, 2016). Microglia alterations were found in several regions of the maternal brain, and normalized by postpartum day 21 in all regions except the hippocampus (Haim et al., 2017). This indicates that changes in the neuroimmune environment of the maternal hippocampus may be longer lasting. The hippocampus undergoes considerable plasticity in the peripartum period (Galea et al., 2014), perhaps not surprisingly given its role in cognitive function (Sweatt, 2004) and mood regulation (Campbell and MacQueen, 2004). Neuroimmune processes are implicated in cognition (Lee et al., 2008; Parkhurst et al., 2013; vom Berg et al., 2012), stress (Hodes et al., 2014; Kreisel et al., 2014), and mood (Setiawan et al., 2015), raising the possibility that changes in the neuroimmune environment of the hippocampus may represent a substrate for motherhood-related changes in hippocampal function.

In the non-maternal brain, immune processes have been implicated in the regulation of adult hippocampal neurogenesis, under basal and inflammatory conditions (reviewed in Sierra et al., 2014). For example, inflammation was first demonstrated to suppress neurogenesis by studies utilizing systemic or intrahippocampal administration of the bacterial endotoxin lipopolysaccharide (Ekdahl et al., 2003; Monje, 2003). Microglia also maintain homeostasis in the healthy adult neurogenic niche via phagocytosis of apoptotic new cells (Sierra et al., 2010). In the maternal brain, one study found that alterations in T-cell activity accounted for at least some of the postpartum-associated reductions in neurogenesis (Rolls et al., 2008). To date, however, no studies have concurrently examined adaptations in microglia and neurogenesis in the maternal hippocampus, and therefore the experiments reported here aimed to fill this gap.

Given the extensive cross-talk between the central nervous system and the immune system (Louveau et al., 2015), motherhood-related adaptations in the immune system can potentially drive plasticity in the brain. Reproductive immunology research has been primarily focused on aspects of immune function that affect fetal development and the success of pregnancy (PrabhuDas et al., 2015). Although many of the maternal immune adaptations normalize in the postpartum period (Groer et al., 2015), there is evidence indicating that maternal experience can leave a lasting footprint on the immune system (Barrat et al., 1997a, 1997b, 1999; Helle et al., 2004). For example, the risk of dying of infectious disease after the age of 65 was increased in mothers of twins, compared to mothers of singletons (Helle et al., 2004). This effect may be related to reproductive effort, and is perhaps indicative of accelerated immunosenescence (Helle et al., 2004). Although the long-term effects of parity on the immune system have received little attention in animal models, delayed senescence in certain aspects of immune function is evidenced in parous relative to non-parous mice (Barrat et al., 1997b, 1997a, 1999). In tandem with neurogenic and neuroimmune markers, our current study aimed to assess whether maternal experience can alter the circulating cytokine profile at various intervals following parturition, ending well after the reproductive event itself. The circulating cytokine profile is not only informative to the general inflammatory state, but may have ramifications for brain and behaviour as peripheral cytokine signals propagate to the brain (Miller et al., 2014; Quan and Banks, 2007). Preclinical cytokine data may also be valuable for comparative purposes, as circulating cytokines levels are accessible biomarkers in clinical populations (Guerreiro et al., 2007).

In this study, we examined the short- and long-term effects of parity on microglia density and morphology, and on neurogenesis in the hippocampus. These measures were examined across age and time since parturition, extending into middle age. To gain information about the peripheral inflammatory milieu, we quantified concentrations of various serum cytokines across the same time points. We expected parity to suppress hippocampal neurogenesis in the short term, and to increase neurogenesis in middle age. At least in the short term, we expected parity to modify microglial density and morphology in the dentate gyrus. Finally, we expected alterations in the circulating cytokine profile during pregnancy and the early postpartum period, and hypothesized that parity would modulate the age-related changes in the circulating cytokine milieu.

## 2. Materials and Methods

### 2.1. Animals

Young adult female and male Sprague-Dawley rats were purchased from Charles River Laboratories (Montreal, Canada), weighing at 200–250g. All rats arrived at our facility at the same time. Rats were maintained on a 12-hour light/dark cycle (lights on at 07:00 h), in standard laboratory conditions (21 ± 1°C; 50 ± 10% humidity) and given *ad libitum* access to water and food (Purina Rat Chow). Female rats were initially pair-housed, and except for the breeding period, all rats were housed in female-only colony rooms. Males were used for breeding purposes only. Nulliparous rats were never housed in the same colony room as primiparous rats when they were breeding or had active litters. To minimize potential environmental exposure differences between nulliparous and primiparous groups, primiparous rats were transferred to the nulliparous colony room on the day that their litters were weaned (postpartum day 21). All procedures were performed in accordance with ethical guidelines set by the Canadian Council on Animal Care and approved by the Animal Care Committee at the University of British Columbia.

### 2.2. Breeding procedure and experimental groups

Female rats were bred at approximately 7 months of age. At 18:00 h daily, each pair of female cage-mates was placed with one male. Vaginal lavage samples were obtained the following morning, between 08:00 and 09:00 h, and examined for the presence of sperm cells. The detection of sperm cells indicated Gestation Day 1 (GD1), at which point the pregnant female was weighed and single-housed. Primigravid or primiparous rats (i.e. pregnant or mothering for the first time; n=30) were randomly assigned to one of six groups (n=5 each) according to the timeline of euthanasia relative to gestation. This included one gestational group at Gestation Day 13 (GD13), and four postpartum groups: Postpartum Day 8 (PPD8), Postpartum Day 30 (PPD30), Postpartum Day 90 (PPD90), and Postpartum Day 180 (PPD180). Nulliparous rats (i.e. never pregnant; n=30) were randomly assigned to control groups (n=5 each) that were age-matched to each of the primiparous groups. Specifically, nulliparous rats at approximately 7, 7.5, 8, 10, and 13 months of age served as control groups for primiparous rats at GD13, PPD8, PPD30, PPD90, and PPD180, respectively (experimental groups are detailed in **Figure 1)**. These timepoints were chosen to capture: mid-gestation (GD13), as a previous study found reductions in the survival of hippocampal neurons produced at this time (Rolls et al., 2008); an early postpartum timepoint (PPD8) that avoids the acute inflammatory state surrounding parturition (Catalano et al., 2010) and is associated with declines in cell proliferation (Leuner et al., 2007); a post-weaning, late postpartum timepoint (PPD30) shown to be associated with reduced neurogenesis (Workman et al., 2015); and finally, for a time-course analysis of the effects of parity on the ageing trajectory, two further timepoints were selected leading to middle age (PPD180), as previous studies reported increased neurogenesis in middle-aged rats with previous maternal experience (Barha et al., 2015; Galea et al., 2018). To account for potential effects of social housing, nulliparous controls were single-housed at GD1 of their primiparous-counterparts and until experimental endpoint. For all postpartum groups, litters were culled to include between 8-10 pups, with approximately 50% males and females. When the original sex ratio or litter size was not sufficient to achieve this, pups were cross-fostered between dams that had given birth the same day. Pups were weaned at PPD21 for all postpartum groups, except for the PPD8 group in which the dams remained with their litter until just prior to euthanasia. After weaning, primiparous rats remained single-housed until experimental endpoint. One rat from the nulliparous 10-month-old group was eliminated from the study due to a mammary gland tumor.

**Figure 1.**
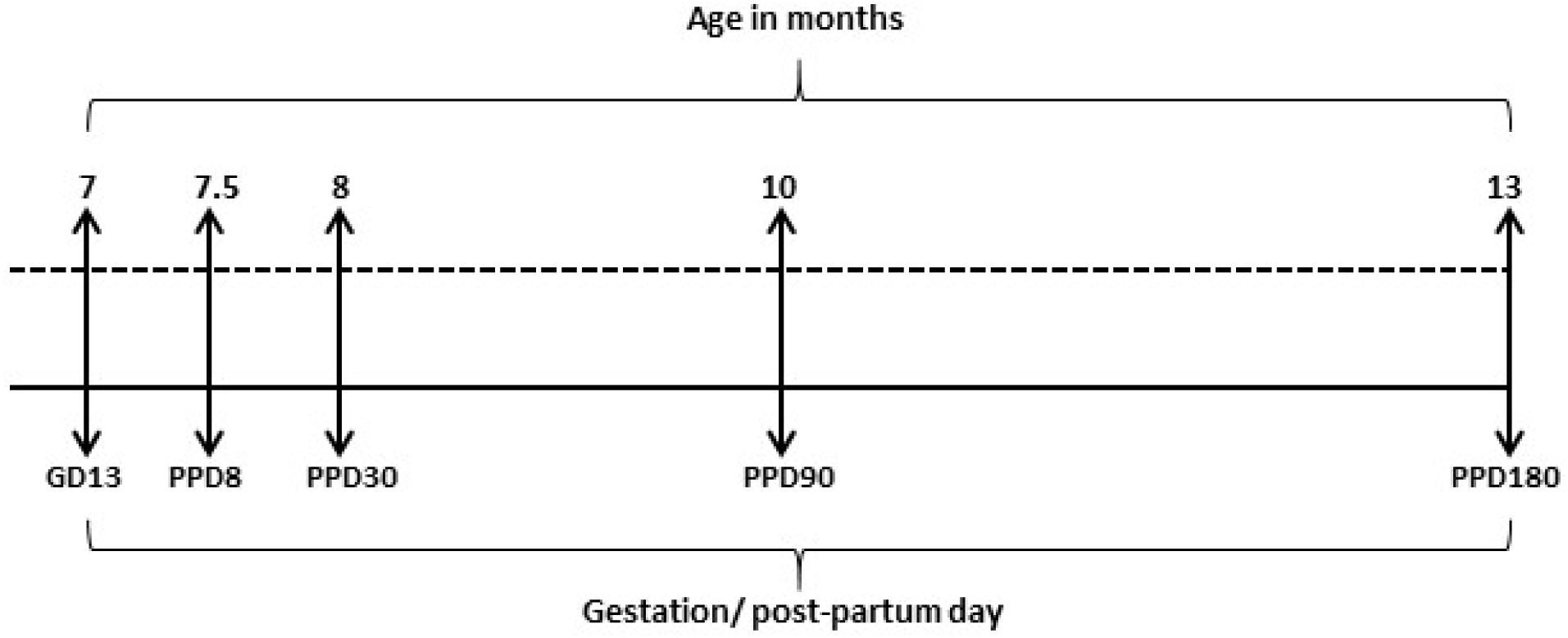
Experimental groups. Primigravid or primiparous rats (n=5/group) were euthanized at gestation day 13 (GD13), postpartum day 8 (PPD8), postpartum day 30 (PPD30), postpartum day 90 (PPD90), or postpartum day 180 (PPD180). Nulliparous rats (n=5/group) were age-matched to their primiparous counterparts, and euthanized along the same timeline, depicted as approximate age in months. Arrows indicate euthanasia day.

### 2.3. Perfusion and tissue collection

All perfusions were completed between 9:00 and 11:00 am. The rats were deeply anesthetized with an overdose of sodium pentobarbital (i.p.), and blood was collected via cardiac puncture. Brains were collected immediately after transcardial perfusion with 60ml of cold 0.9% saline, followed by 120ml of cold 4% paraformaldehyde (PFA). Brains were stored at 4°C in 4% PFA for 24 hours, then transferred into a 30% sucrose solution (in 0.1 M Phosphate Buffer) until sectioning. In the group euthanized during gestation, the uterus was dissected to confirm pregnancy. To obtain serum, blood samples were allowed to clot overnight at 4°C, then centrifuged at 10g for 15 minutes and serum aliquots were stored at −20°C until processing.

### 2.4. Brain Tissue Processing and Immunohistochemistry

Brains were sliced into 40 µm coronal sections using a Leica SM2000R Microtome (Richmond Hill, Ontario, Canada). Sections were collected in series of 10 along the rostral-caudal axis of the hippocampus, then stored at −20 °C in a cryoprotectant consisting of 30% ethylene glycol (Sigma-Aldrich, St. Louis, MO, USA) and 20% glycerol (Sigma-Aldrich) in 0.1 M phosphate-buffer (PB, pH 7.4). Sections were thoroughly rinsed (5 x 10 mins) in PBS prior to staining to remove the cryoprotective medium. All immunohistochemical procedures were conducted on free-floating brain sections, and on a rotator at room temperature unless otherwise noted.

#### 2.4.1 Doublecortin (DCX)

DCX is a microtubule-associated protein expressed in immature neurons for 21 days after production in adult rats (Brown et al., 2003), and thus was used as a marker of adult hippocampal neurogenesis. Tissue was rinsed in 0.1M PBS (pH 7.4; 5 x 10 minutes) between each of the following procedures. Tissue was treated with 0.3% hydrogen peroxide (H_2_O_2_, in dH_2_O) for 30 minutes, then incubated for 24 hours at 4°C in a primary antibody solution containing 1:1000 goat anti-doublecortin (Santa Cruz Biotechnology, Santa Cruz, CA, USA) in 3% normal rabbit serum (NRS) and 0.4% Triton-X in 0.1M PBS. Next, tissue was transferred to a secondary antibody solution consisting of 1:500 rabbit anti-goat (Vector Laboratories, Burlington, ON, Canada) in 0.1 M PBS for 24 hours at 4°C. Finally, tissue was transferred to an avidin-biotin complex (ABC; Elite kit; 1:1000, Vector Laboratories) in PBS for 4 hours, then immunoreactants were visualized with a Nickel-enhanced DAB reaction (Vector Laboratories). Sections were mounted onto glass slides and allowed to dry, then dehydrated in increasing graded ethanol, defatted with xylenes, and cover-slipped with Permount (Fisher Scientific).

#### 2.4.2 Ionized calcium binding adaptor molecule-1 (Iba-1)

Iba-1 is a calcium-binding protein widely used as a microglial marker (Korzhevskii and Kirik, 2016). Tissue was rinsed in 0.1 M PBS (pH 7.4; 3 x 10 minutes) between each of the following procedures. Tissue was incubated in 0.3% hydrogen peroxide (H_2_O_2_, in dH_2_O) for 25 minutes, then blocked with 10% normal goat serum (NGS) in 0.5% Triton-X in 0.1M PBS. Tissue was then transferred to a primary antibody solution for 18 hours at 4°C, consisting of 1:1000 anti-Iba-1 (Wako, Osaka, Japan) in 10% NGS and 0.4% Triton-X in 0.1M PBS. Next, tissue was incubated in a secondary antibody solution for 1 hour, containing 1:500 biotinylated anti-rabbit (Vector Laboratories) in 2.5% NGS and 0.4% Triton X in PBS. Finally, tissue was transferred to an avidin-biotin complex (ABC; Elite kit; 1:50, Vector Laboratories) in 0.4% Triton-X in PBS for 1 hour, and immunoreactivity was visualized with a Nickel-enhanced DAB reaction (Vector Laboratories). Sections were mounted onto glass slides and allowed to dry, then counterstained with cresyl violet, dehydrated in a series of ethanol solutions of increasing concentrations, defatted with xylenes, and cover-slipped with Permount (Fisher Scientific).

#### 2.4.3 Ki67

Ki67 is expressed during all active phases of the cell cycle, but not during G_0_ phase (Scholzen and Gerdes, 2000), and therefore was used as a marker of cell proliferation in the dentate gyrus. Tissue was rinsed in 0.1 M PBS (pH 7.4; 3 x 10 minutes) between each of the following procedures. Tissue was incubated in a primary antibody solution for 48 hours at 4°C, consisting of 1:200 mouse anti-Ki67 (NCL-L-Ki67-MM1; Leica Biosystems, Newcastle, UK) in 3% normal donkey serum (NDS), and 0.3% Triton-X in 0.1M PBS. Next, tissue was incubated for 18 hours at 4°C in a secondary antibody solution consisting of 1:200 donkey anti-mouse IgG, Alexa Fluor 555 (Molecular Probes, Eugene, Oregon, USA) in 3% NDS in 0.1M PBS. Sections were counterstained with DAPI (2.5 minutes; 300nM; ThermoFisher, Waltham, WA, USA), mounted onto glass slides, and cover-slipped with an anti-fade medium (2.5% Polyvinyl alcohol-Dabco).

### 2.5. Microscopy, Cell Quantification, and Morphological analyses

An investigator blinded to experimental conditions quantified DCX- and Iba-1-, and Ki67-immunoreactive cells and analyzed cell morphology. See **Fig. 4B, 6D-F, and 7** for representative photomicrographs.

#### 2.5.1 Iba-1

Under a 40x objective on a Nikon E600 microscope, an exhaustive quantification of Iba-1-IR cells was completed in four hippocampal slices from each animal, as we have done previously (Mahmoud et al., 2016a). This included two dorsal and two ventral sections, with approximate Bregma coordinates of −3.12, −3.48, 6.00, and −6.36. Iba-1-IR cells were quantified in the dentate gyrus, specifically in the granule cell layer (GCL), the subgranular zone (SGZ), and within an approximately 50μm band of the molecular layer (ML).

Under baseline conditions microglia typically display ramified morphology, characterized by highly branched and long processes that continuously survey their environment (Nimmerjahn et al., 2005). Under inflammatory states, microglial processes typically retract and the cell can take on an amoeboid morphology, in which the cell become enlarged and void of processes; thus, microglial morphology is widely used as a proxy-measure of functional state (Karperien et al., 2013). Here, Iba-1-IR cell morphology was analyzed utilizing NIS Elements Basic Research software (Nikon) under a Nikon E600 microscope. Using the measure feature, soma size, in addition to cell process length and number (Karperien et al., 2013) were measured live at 40x for every cell within a 23672.24 μm^2^ region of interest (ROI), with 3 ROIs each for the dorsal and ventral hippocampus. Further, no more than one ROI was taken from an individual tissue slice, and ROIs were defined in 3 consistent locations in the GCL for both the dorsal and ventral hippocampus within each animal. Cells were defined by the presence of an Iba-1-IR cell body within the ROI, and this definition did not necessitate the presence of cell processes, by that ensuring the inclusion of any cells with amoeboid morphology. The average process length per cell was calculated using the total length and number of processes for each cell, and subsequently an average process length was calculated for each animal. Both primary (extending directly from the cell body) and secondary processes were taken into account in the analyses. There were no significant differences between groups in the number of Iba-1-IR cells that fell within the ROIs and were used for morphological analyses (see Table 1).

**Table 1.** Mean number of Iba-1-IR cells used for process length analyses ± standard error of the mean. There were no significant differences between groups.

| Group | Number of Iba-1-IR cells analyzed |
| --- | --- |
| Nulliparous – 7 mo. | 23.40 ± 1.03 |
| Nulliparous – 7.5 mo. | 21.00 ± 2.04 |
| Nulliparous – 8 mo. | 20.80 ± 2.13 |
| Nulliparous – 10 mo. | 21.20 ± 1.16 |
| Nulliparous – 13 mo. | 22.25 ± 2.32 |
| Primiparous – GD13 | 23.67 ± 1.67 |
| Primiparous – PPD8 | 22.00 ± 0.77 |
| Primiparous – PPD30 | 23.60 ± 1.47 |
| Primiparous – PPD90 | 20.60 ± 1.03 |
| Primiparous – PPD180 | 23.20 ± 0.37 |

In an additional analysis of microglial morphology, Iba-1-IR cells were categorized into one of three morphological phenotypes, adapted from other methods (Haim et al., 2017; Schwarz et al., 2012). Specifically, as depicted in the representative photomicrographs in Fig. **6D-F**, cells were categorized as 1) ramified microglia (highly branched, with longer processes), stout microglia (few, shorter processes), or 3) amoeboid microglia (no processes). This was performed under a 40X objective, in 4 hippocampal slices per animal, and involved a classification of all Iba-1-IR cells in the GCL, SGZ, and an approximately 50μm band of the ML. For each animal, the percentage of cells of each morphological phenotype was calculated.

#### 2.5.2 DCX

Under a 100x objective on an Olympus CX22LED brightfield microscope, DCX-IR cells in the granule cell layer (GCL) were exhaustively counted in every 10^th^ section of the hippocampus along the rostral-caudal axis. Thus, raw counts were multiplied by a factor of 10 to obtain an estimate of the total number of DCX-IR cells in the hippocampus. Previous work indicates that certain conditions or experiences, including reproductive experience (Workman et al., 2015), can alter the rate at which newly produced hippocampal neurons mature (Overstreet-Wadiche et al., 2006), and this can have functional consequences as changes in the maturational timeline can alter the rate of functional integration of these new neurons into existing hippocampal networks. Thus, to examine whether pregnancy and motherhood can alter the rate at which newly produced hippocampal neurons mature, we investigated the maturational stages of DCX-immunoreactive cells. Specifically, using the 100× objective on an Olympus CX22LED brightfield microscope, 50 DCX-IR cells (25 dorsal GCL and 25 ventral GCL; each taken from 3 slices) were randomly selected for each animal for maturational staging. Cells were categorized into one of three maturational stages, based on previously established criteria (Plümpe et al., 2006): 1) proliferative (no process or short process), 2) intermediate (medium process with no branching), or 3) post-mitotic (long processes with branching in the GCL and ML). DCX-immunolabeled sections were also utilized to measure GCL areas in every 10^th^ section of the hippocampus, using images taken at 4x and the ImageJ software. Volume was estimated using these areas and following Cavalieri’s principle.

#### 2.5.3 Ki67

Under a 100x objective on an Olympus CX22LED microscope equipped with epifluorescence, Ki67-IR cells in the GCL and SGZ of the DG were exhaustively counted in every 10^th^ section of the hippocampus along the rostral-caudal axis. Raw counts were multiplied by a factor of 10 to obtain an estimate of the total number of Ki67-IR cells in the DG.

### 2.6. Serum cytokine and hormone quantification

A multiplex immunoassay kit (V-PLEX Proinflammatory Panel 2, Rat) was purchased from Meso-Scale Discovery (Rockville, MD) and used according to manufacturer instructions to measure serum cytokine levels. The antibody pre-coated plates allowed for the simultaneous quantification of the following cytokines: Interferon-gamma (IFN-γ), Interleukin-1beta (IL-1β), Interleukin-4 (IL-4), Interleukin-5 (IL-5), Interleukin-6 (IL-6), chemokine (C-X-C motif) ligand 1 (CXCL1), Interleukin-10 (IL-10), Interleukin-13 (IL-13), and tumor necrosis factor alpha (TNF-α). Samples were run in duplicates, and plates were read with a Sector Imager 2400 (Meso Scale Discovery), and data was analyzed using the Discovery Workbench 4.0 software (Meso Scale Discovery). The assays’ lower limits of detection (LLOD), which varied between analytes and plates (2 plates total), were as follows (pg/mL): IFN-γ: 0.163-0.266; IL-10: 0.233-0.313; IL-13: 0.78-2.7; IL-1β:1.48-1.62; IL-4:0.179-0.298; IL-5: 7.64-9.8; IL-6: 2.48-2.49; CXCL1: 0.085-0.164; and TNF-α: 0.156–0.186. Any values below the LLOD were assigned 0 pg/mL, as we have done previously (Bodnar et al., 2017). All samples were within the detection range for TNF-α, CXCL1, and IL-10. One sample fell below the LLOD for each of IFN-γ, IL-4, and IL-13. For three cytokines, a number of samples fell below the LLOD (n=12 for IL-6, n=17 for IL-1β, and n=36 for IL-5). This panel was chosen as it includes a broad range of cytokines, some traditionally considered proinflammatory (IL-1β, IFN-γ, TNF-α), anti-inflammatory (IL-4, IL-10), and pleiotropic (IL-6), in addition to the chemokine CXCL1 which is important for neurotrophil recruitment. Therefore, combined, these markers provide a comprehensive view of the inflammatory milieu.

Steroids hormones modulate neuroplasticity (Mahmoud et al., 2016b) and interact with the immune system (Schumacher et al., 2014). Further, circulating concentrations increase dramatically during pregnancy, drop abruptly at parturition, and diminish in ageing females. To investigate the potential mediating roles of steroid hormones, we measured 17β-estradiol, the most potent of the estrogens and the most abundant in premenopausal women and younger female rats (Rannevik et al., 1986). Serum 17β-estradiol concentrations were quantified in sample taken at perfusion, using an ultra-sensitive estradiol radioimmunoassay (Beckman Coulter, Prague, Czech Republic) and in accordance with manufacturer’s instructions, with all samples run in duplicates. The assay sensitivity is 2.2pg/ml, and the average inter- and intra-assay coefficients of variation were <15%.

### 2.7. Statistical analyses

Statistical analyses were performed using Statistica software (Tulsa, OK). All dependent variables were subjected to the Kolmogorov-Smirnov test for normality, and Bartlett’s test for heterogeneity of variance. TNF-α, IL-1β, IL-5, and IL-6 data were not normally distributed and therefore Box-Cox transformed prior to analyses. There were few violations to the heterogeneity of variance assumption, however these were corrected after Box-Cox transformation. Neural measures (DCX- and Ki67-IR cell number, Iba-1-IR density, length and number of processes) and serum measures (17β-estradiol, IFN-γ, IL-1β, IL-4, IL-5, IL-6, CXCL1, IL-10, IL-13, and TNF-α) were each analyzed using factorial analysis of variance (ANOVA), with time (GD13/7 mo., PPD8/7.5 mo., PPD30/8 mo., PPD90/10 mo., PPD180/13 mo.) and reproductive status (primi-gravid/parous, nulliparous) as the between-subject factors. Post-hoc analyses utilized Fisher’s LSD. A priori we expected parity to modulate the age-related changes in cytokine levels, and density/morphology of Iba-1-IR cells. Any a priori comparisons were subjected to a Bonferroni correction. Pearson’s correlations were performed on dependent variables of interest, and Fisher’s z-transformation of correlation coefficients was used to assess the significance of the difference between correlations in nulliparous and primiparous groups. Serum 17β-estradiol concentrations were used as a covariate in ANOVAs, as circulating concentrations can influence neuroplastic and immune measures (Mahmoud et al., 2016b; Schumacher et al., 2014); ANOVAs are reported with the covariate in instances where there was a significant main effect of the covariate. Finally, as an exploratory approach, and to complement our findings from ANOVA analyses, we ran a principal component analysis (PCA) on the cytokine data, with the purpose of deriving information about the amount of variance accounted for by potential cytokine networks within the dataset.

## 3. Results

### 3.1. Serum 17β-estradiol was reduced in the early postpartum and increased in the late postpartum

As expected, 17β-estradiol was reduced in the early postpartum period at PPD8, relative to age-matched nulliparous rats (p=0.015; reproductive status by time interaction: F(4, 36)=4.45, p=0.005; **Fig. 2**). Further, primiparous rats at PPD30 had higher serum 17β-estradiol concentrations than age-matched nulliparous rats (p<0.008; **Fig. 2**). Finally, 17β-estradiol concentrations at PPD30 and PPD90 were significantly higher than all other primiparous groups (p’s < 0.03), but there were no significant differences between any of the nulliparous groups (all p’s >0.19). There was also a significant main effect of time (p<0.01) but not of reproductive status (p=0.41).

**Figure 2.**
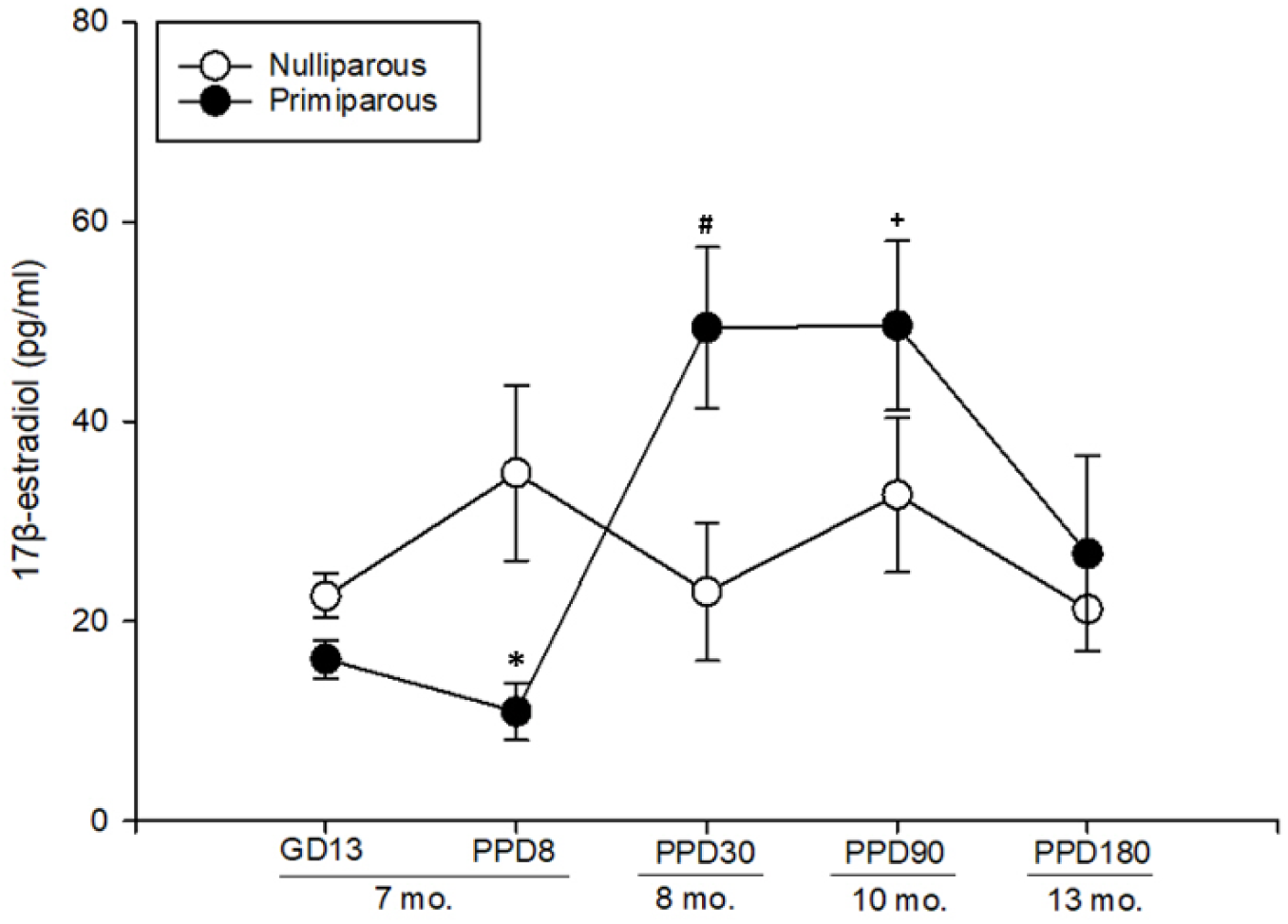
Serum concentrations of 17β-estradiol at perfusion in primiparous and nulliparous rats. The x-axis represents time relative to gestation and parturition in primiparous rats, and approximate age in months. 17β-estradiol was significantly reduced in the early postpartum period at PPD8, but significantly increased in the late postpartum period at PPD30 and PPD90. * indicates p = 0.015, significantly lower than age-matched nulliparous rats; # indicates p <0.03, significantly higher that 8-month-old nulliparous rats, and primigravid/parous rats at GD13, PPD8, and PPD180; + indicates p <0.03, significantly higher than primigravid/parous rats at GD13, PPD8, and PPD180. Data are represented in mean values ± SEM. GD= gestation day, PPD= postpartum day.

### 3.2. Parity and age had no significant effect on granule cell layer volume

Granule cell layer volume was not significantly affected by age, parity, or age by parity interaction (all p’s >0.48), thus all further analyses were performed on the number, rather than density, of Ki67- and DCX-IR cells.

### 3.3. The number of doublecortin-IR cells was significantly reduced during pregnancy and the postpartum period, and declined in middle-age in nulliparous rats only

The number of DCX-IR cells was significantly reduced in primi-gravid and -parous rats relative to age-matched nulliparous controls at GD13, PPD8, and PPD30 (p’s <0.001; significant time by reproductive status interaction; F(4, 38)=6.7213, p<0.001; **Fig. 3**). There were also significant main effects of time and reproductive status (all p’s < 0.001).

**Figure 3.**
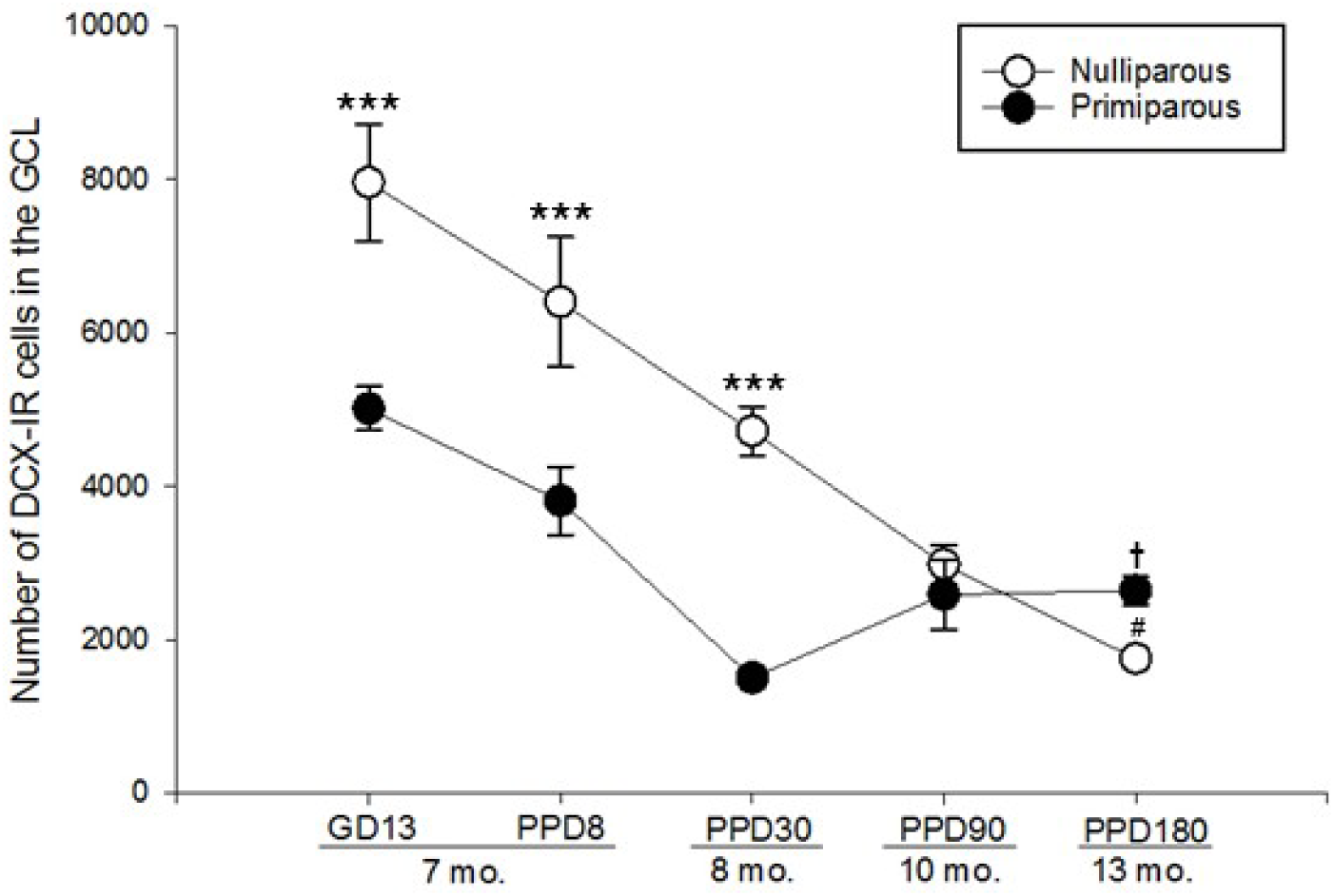
Estimated total number of doublecortin (DCX)-immunoreactive (IR) cells in the granule cell layer of primiparous and nulliparous rats across 7-13 months of age. The x-axis represents time relative to gestation and parturition in primiparous groups, and approximate age in months. DCX-IR cell number was significantly reduced in primiparous rats in mid-gestation and until postpartum day 30. Between 8 and 13 months of age, DCX-IR cell number significantly declined in nulliparous rats, but significantly increased in primiparous rats. Data are represented in mean values ± standard error of the mean (SEM). GCL= granule cell layer, DCX-IR= doublecortin-immunoreactive, GD= Gestations Day, PPD= Postpartum Day, mo.= approximate age in months. *** indicates p<0.001, significantly different from age-matched primiparous group. # denotes p<0.001, significantly different from 8-month-old nulliparous group. † indicates p=0.044, significantly different from primiparous rats at PPD30.

There was a significant age-related decline in the number of DCX-IR cells in nulliparous rats, such that each nulliparous group had a significantly lower number of DCX-IR cells than all previous nulliparous age groups (all p’s<0.014), with the exception of a non-significant decline from 10 to 13 months (p=0.11; significant time by reproductive status interaction; F(4, 38)=6.7213, p<0.001; **Fig. 3**). There was also a significant decline in DCX-IR cell number in primiparous rats at PPD30 relative to GD13 and PPD8 (p’s<0.017; **Fig. 3**). However, the trajectory of age-related change in DCX-IR cell number was significantly altered by parity in middle-age; unlike nulliparous rats, primiparous groups had no significant difference in the number of DCX-IR cells from 8 to 10 months of age (p=0.10). Based on previous findings (Barha et al., 2015; Galea et al., 2018), we expected parity to increase neurogenesis in middle age. Indeed, we found a significant increase in DCX-IR cells from 8 to 13 months of age in primiparous rats (p=0.044; one-tailed).

### 3.4. Parity and age had no significant effect on the maturational stage of DCX-IR cells

Regardless of age and reproductive status, the percentage of proliferative DCX-IR cells was significantly higher than that of intermediate DCX-IR cells (p<0.001), and the percentage of post-mitotic DCX-IR cells was significantly higher than that of intermediate and proliferative DCX-IR cells (p’s<0.001; significant main effect of DCX maturational stage, F(2, 76)=107.17, p<0.001; **Table 2**).

**Table 2.** Mean percentage of proliferative, intermediate, and post-mitotic doublecortin (DCX)-immunoreactive (IR) cells in the granule cell layer ± standard error of the mean. Parity and age did not significantly affect the maturational stage of DCX-IR cells.

| Group | % Proliferative | % Intermediate | % Post-mitotic |
| --- | --- | --- | --- |
| Nulliparous – 7 mo. | 37.60 ± 1.60 | 17.20 ± 2.15 | 45.20 ± 2.65 |
| Nulliparous – 7.5 mo. | 26.40 ± 2.86 | 18.80 ± 1.85 | 54.80 ± 2.87 |
| Nulliparous – 8 mo. | 35.60 ± 4.71 | 20.80 ± 1.02 | 46.40 ± 4.45 |
| Nulliparous – 10 mo. | 29.60 ± 6.52 | 21.60 ± 2.64 | 48.80 ± 4.84 |
| Nulliparous – 13 mo. | 25.33 ± 8.74 | 20.00 ± 1.15 | 54.67 ± 8.97 |
| Primiparous – GD13 | 32.00 ± 6.23 | 18.40 ± 2.93 | 49.60 ± 5.84 |
| Primiparous – PPD8 | 28.40 ± 5.31 | 16.00 ± 1.10 | 56.40 ± 6.71 |
| Primiparous – PPD30 | 27.60 ± 4.26 | 17.20 ± 0.80 | 55.20 ± 4.88 |
| Primiparous – PPD90 | 23.60 ± 3.19 | 16.80 ± 3.01 | 59.60 ± 4.53 |
| Primiparous – PPD180 | 28.40 ± 5.71 | 16.00 ± 2.45 | 56.00 ± 6.23 |

There were no other significant main effects, and no significant interactions (all p’s >0.14).

### 3.5. Ki67-IR cells declined in mid-gestion and the early postpartum period in primiparous rats, and with age regardless of reproductive status

Parity significantly reduced the number of Ki67-IR cells in the GCL (F(1, 38)=9.2681, p=0.004; significant main effect of reproductive status; **Fig. 4A**). Regardless of reproductive status, Ki67-IR cells declined significantly with age (F(4, 38)=16.505, p<0.001; main effect of time; **Fig. 4A**), in which there was a significant difference between all age groups (p’s < 0.005), with the exception of non-significant differences from 7 to 7.5 months, 8 to 10 months, and 10 to 13 months (p’s > 0.05). Although there was no significant age by parity interaction (p=0.13), a priori we expected a decline in cell proliferation in the early postpartum period based on previous work (Leuner et al., 2007). Planned comparisons revealed a significant decline in Ki67-IR cells on PPD8 (p=0.013; one-tailed), in addition to a significant decline on GD13 (p= 0.004), relative to age-matched nulliparous controls in both instances (**Fig. 4A**).

**Figure 4.**
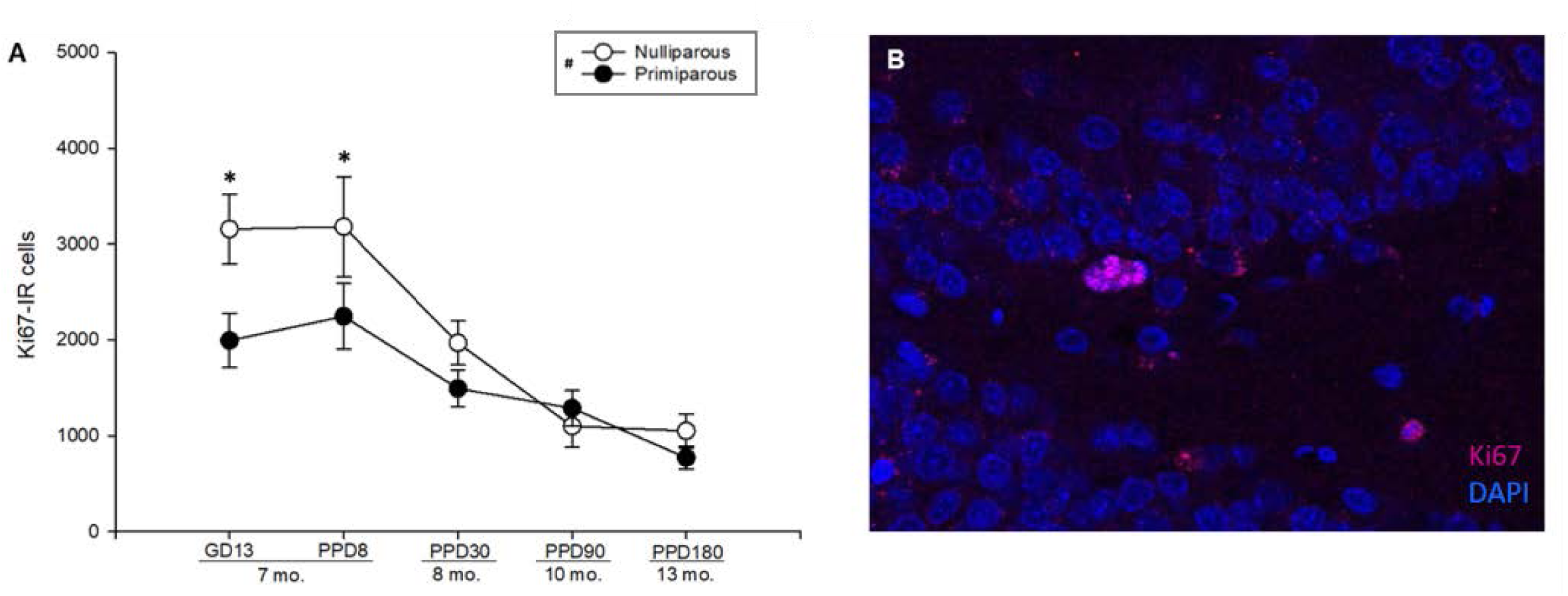
**(A)** Estimated total number of Ki67-immunoreactive (IR) cells in the granule cell layer and subgranular zone of primiparous and nulliparous rats across 7-13 months of age. The x-axis represents time relative to gestation and parturition in primiparous groups, and approximate age in months. Ki67-IR cell number was significantly reduced in primiparous rats in mid-gestation and the early postpartum period, and declined with age regardless of reproductive status. Data are represented in mean values ± standard error of the mean (SEM). Ki67-IR = Ki67-immunoreactive, GD= Gestations Day, PPD= Postpartum Day, mo.= approximate age in months. * indicates p<0.014, significantly different from age-matched primiparous group. # denotes p=0.004, significant main effect of reproductive status. **(B)** Representative photomicrograph of the dentate gyrus, showing Ki67-IR cells (pink), counterstained with DAPI (blue).

### 3.6. Density of Iba-1-IR cells increased in the late postpartum period and fluctuated significantly with age in primiparous but not in nulliparous rats

Because we quantified Iba-1-IR cells in 4 sections per animal, the density of IR cells was analyzed, as we have done previously (Mahmoud et al., 2016a). Age significantly affected the density of Iba-1-IR cells in the dentate gyrus, with higher density at 8 and 13 months relative to 7.5 and 10 months of age (p’s<0.02; significant main effect of time, F(4, 39)=3.43, p=0.017). We expected alterations in the density of Iba-1-IR cells in the postpartum period due to previous findings (Haim et al., 2017). A priori comparisons show that the density of Iba-1-IR cells fluctuated significantly across time in primiparous rats; density was increased in the late postpartum period at PPD30 relative to PPD8 (p=0.002), and in middle-aged rats at PPD180 relative to PPD8 (p<0.003; **Fig. 5A**). On the other hand, density did not significantly change across age in nulliparous rats (all p’s >0.17; **Fig. 5A**). There were no significant differences between nulliparous and primiparous groups across age (all p’s >0.3), except for a trend for higher density at PPD30 relative to age-matched nulliparous controls (p=0.09; **Fig. 5A**). There was no significant main effect of reproductive status and no significant interaction (p’s>0.28).

**Figure 5.**
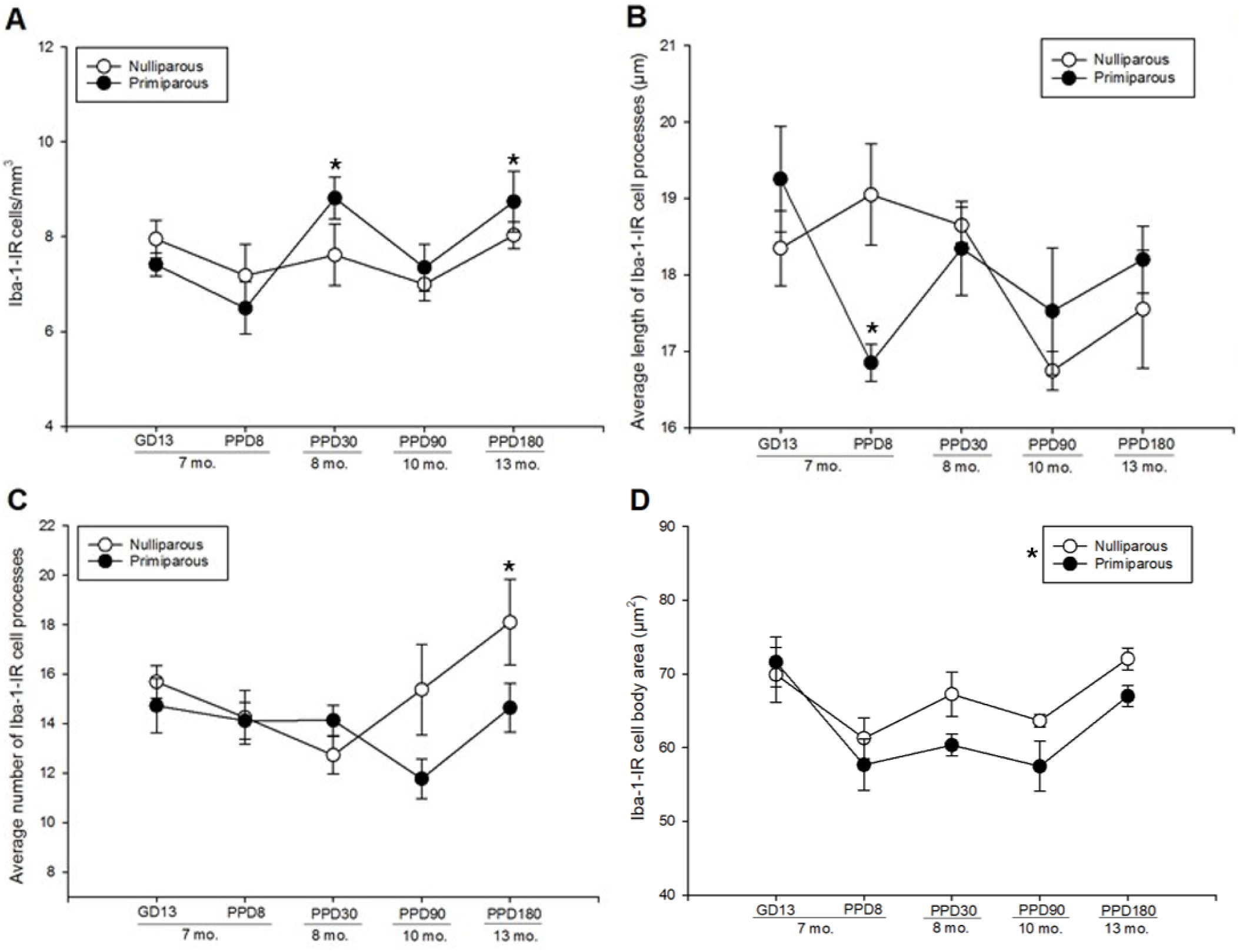
Density and morphology of Iba-1-immunoreactive cells in the dentate gyrus of primiparous and nulliparous rats. The x-axis represents time relative to gestation and parturition in primiparous rats, and approximate age in months. **(A)** Iba-1-IR cell density was stable across age in nulliparous rats, but fluctuated significantly in primiparous rats, with increased density at PPD30 and PPD180, relative to PPD8. * indicates p<0.003, significantly different from primiparous group at PPD8. **(B)** Average length of Iba-1-immunoreactive cell processes was reduced at PPD8, and decreased with age regardless of reproductive status. * denotes p<0.006, significantly different from age-matched nulliparous controls, and from primiparous group at GD13 **(C)** Average number of Iba-1-immunoreactive cell processes increased significantly with age in nulliparous but not primiparous rats. * indicates p<0.002, significantly different from 8-month-old nulliparous group. **(D) There was a trend towards significance for parity to** was reduced soma area of Iba-1-IR cells * indicates p=0.092, main effect of reproductive status. Data are represented in mean values ± SEM. GD= gestation day, PPD= postpartum day, mo.= age in months, Iba-1-IR= Iba-1-immunoreactive.

### 3.7. Average length and number of Iba-1-IR cell processes changed significantly with age and reproductive status

Iba-1-IR cells displayed significantly shorter processes in the early postpartum period at PPD8 relative to nulliparous controls (p=0.005; **Fig. 5B**), and to primiparous rats at GD13 (p<0.002). There was a decline in average length of Iba-1-IR cell processes at 10 months of age, relative to 7 months in primiparous rats, and to 7, 7.5, and 8 months in nulliparous rats (all p’s <0.027; significant time by reproductive status interaction F(4, 38)=4.67, p=0.004; **Fig. 5B**). However, there was no further significant change in average process length between 10 and 13 months of age, regardless of reproductive status (p’s > 0.23). There was also a significant main effect of the covariate serum 17β-estradiol (p = 0.012), a trend towards a significant main effect of time (p=0.067), but no significant main effect of reproductive status (p=0.42).

The average number of Iba-1-IR cell processes increased significantly with age in nulliparous rats, where significantly more processes were found in 13-compared to 8-month-old rats (p<0.002), but missed significance compared to 7.5-month-old rats (p=0.027; planned comparisons; **Fig. 5C**). On the other hand, there were no significant differences in the average number of cell processes between primiparous rats across age (all p’s >0.06). There were trends towards significant main effects of time (p=0.054) and reproductive status (p=0.056) but no significant time by reproductive status interaction (p=0.11).

There was a trend for primiparity to decreased the soma size of Iba-1-IR cells relative to nulliparity (F(1, 38)=5.2646, p=0.092; main effect of reproductive status; **Fig. 5D**). While there was no significant age by parity interaction (p=0.80), the effects of primiparity to reduce soma size appears to be driven by the PPD30, 90, and 180 groups. In addition, there was a significant main effect of time (F(4, 38)=5.28, p<0.001).

### 3.8. The percentage of Iba-1-IR cells of stout morphology increased, and that of ramified morphology decreased, in the early postpartum period

There was a significant Iba-1 morphology by time by reproductive status interaction (F(8, 74)=2.066, p=0.05; **Fig.6**), in which the percentage of ramified Iba-1-IR cells was reduced in primiparous rats at PPD8 relative to age-matched nulliparous controls, primigravid rats at GD13, and primiparous rats at PPD30 (all p’s <0.02; **Fig. 6A**). In addition, the percentage of stout Iba-1-IR cells was significantly increased in primiparous rats at PPD8 relative to age-matched nulliparous controls and primigravid rats at GD13 (p’s <0.007; **Fig.6B**). There were no significant differences in the percentage of amoeboid Iba-1-IR cells between any of the groups (p’s > 0.05; **Fig.6C**). There was also a significant main effect of Iba-1 morphology (p <0.001), a trend for a significant Iba-1 morphology by time interactions (p=0.087), but no other significant main effects or interactions (p’s > 0.49).

**Figure 6.**
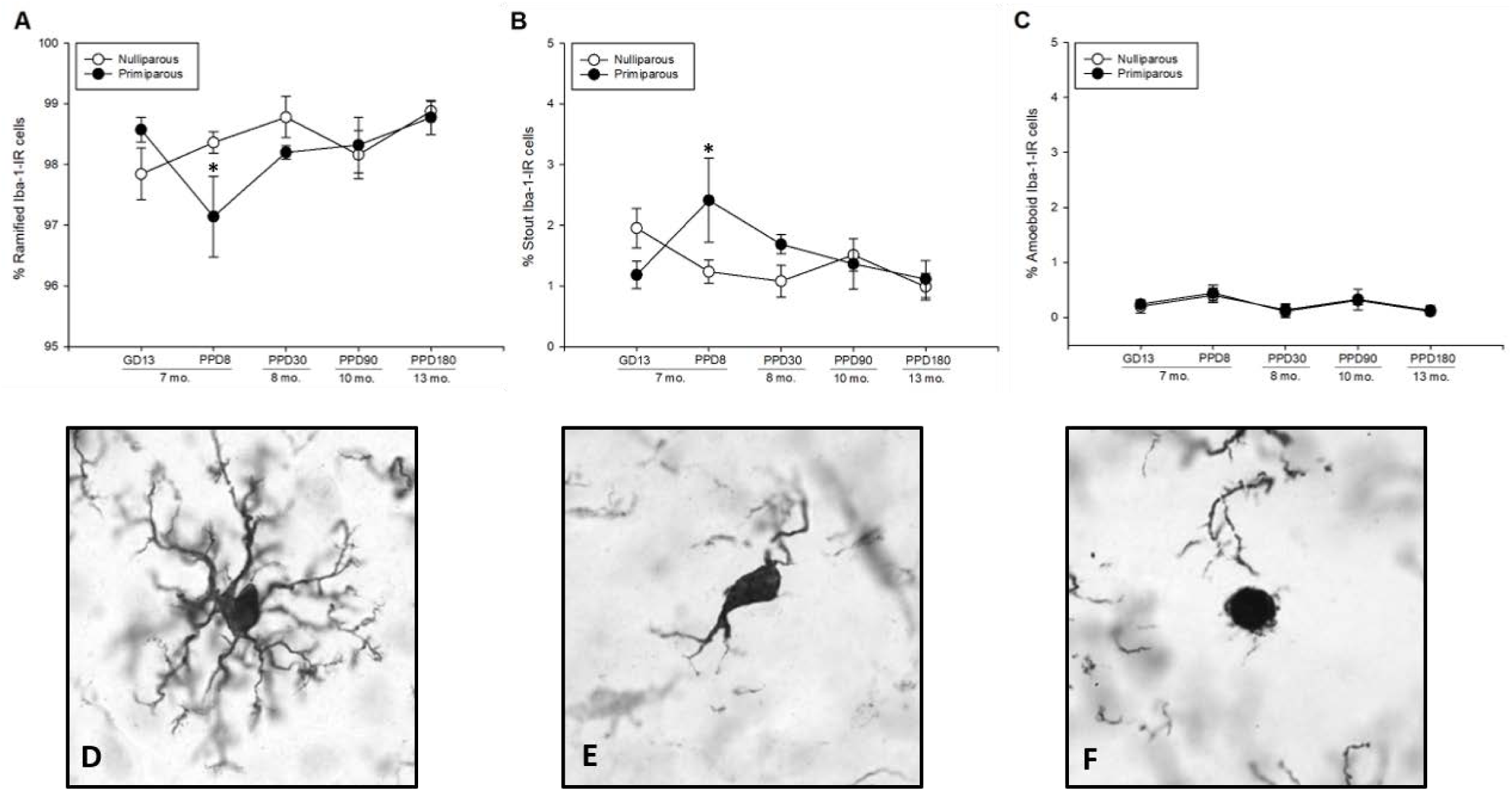
Percentage of Iba-1-immunoreactive cells of ramified **(A)**, stout **(B)**, and amoeboid **(C)** morphology in the dentate gyrus. The x-axis represents time relative to gestation and parturition in primiparous rats, and approximate age in months. **(A)** the percentage of ramified Iba-1-IR cells was transiently reduced in the early postpartum period. * indicates p<0.02, PPD8 significantly lower than 7.5-month-old nulliparous rats, primigravid rats at GD13, and primiparous rats at PPD30. **(B)** the percentage of stout Iba-1-IR cells was transiently increased in the early postpartum period. * indicates p<0.007, PPD8 significantly higher than 7.5-month-old nulliparous rats and primigravid rats at GD13. **(C)** the percentage of amoeboid Iba-1-IR cells did not differ between groups. Representative photomicrographs of Iba-1-IR cells of ramified **(D)**, stout **(E)** and amoeboid **(F)** morphology captured at 60x. Data are represented in mean values ± SEM. GD= gestation day, PPD= postpartum day, mo.= age in months, Iba-1-IR= Iba-1-immunoreactive.

**Figure 7.**
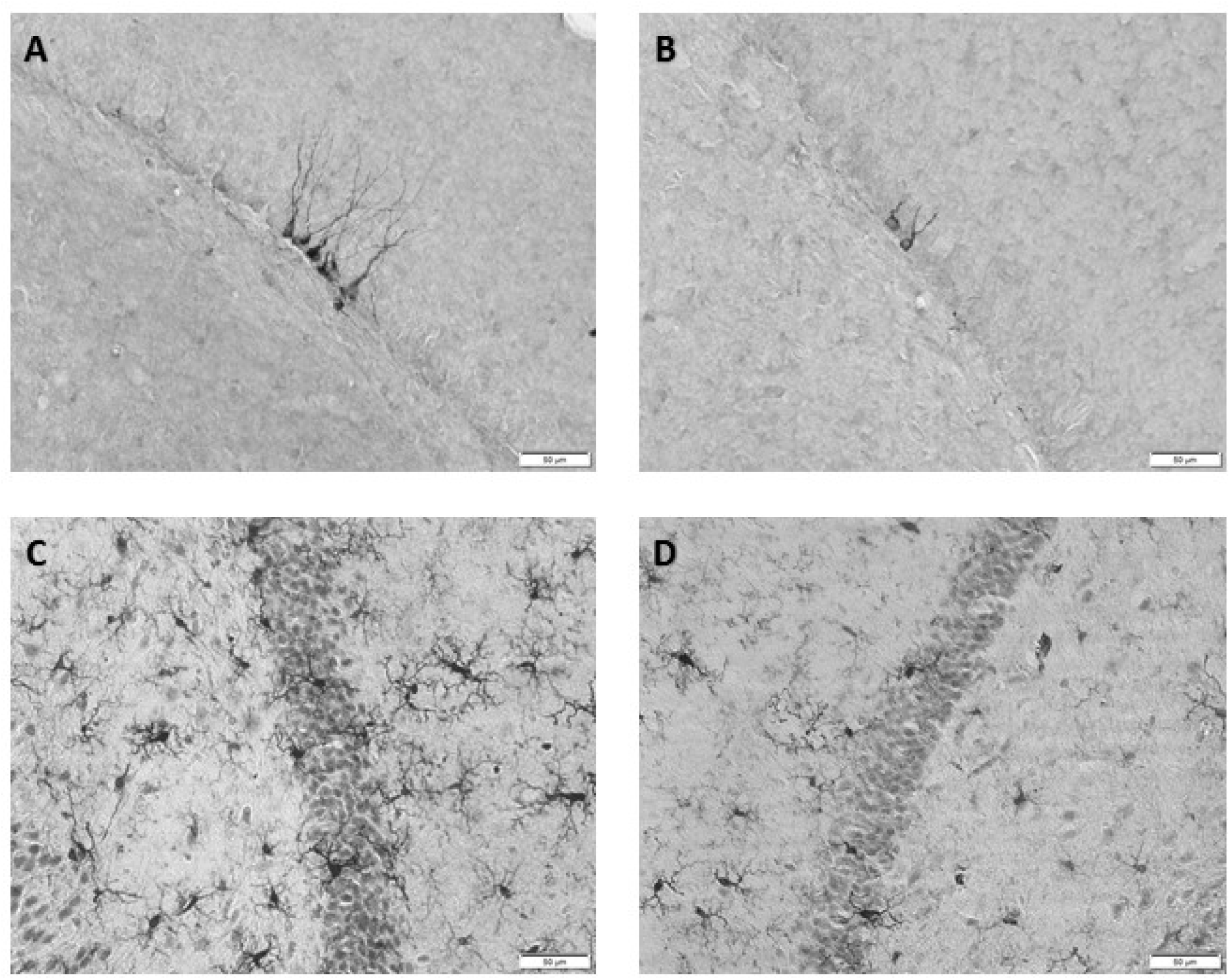
Representative photomicrographs of the granule cell layer in the dentate gyrus. **(A)** Doublecortin-immunoreactive cells in an 8-month-old nulliparous rat. **(B)** Doublecortin-immunoreactive cells in an 8-month-old primiparous rat on postpartum day 30. **(C)** Iba-1-immunoreactive cells in a 7.5-month-old nulliparous rat. **(D)** Iba-1-immunoreactive cells in a 7.5-month-old primiparous rat on postpartum day 8.

### 3.9. Serum IFN-γ and IL-10 showed an age-related increase in nulliparous but not primiparous rats

There was a significant ageing-related increase in IFN-γ in nulliparous rats, in which 13-month-old nulliparous rats had significantly higher IFN-γ levels than all other nulliparous groups (all p’s <0.004; planned comparisons; **Fig. 8A**). No significant differences were found in IFN-γ levels between any primiparous groups (all p’s >0.04, non-significant due to Bonferroni correction). Further, at 13 months, IFN-γ levels were higher in nulliparous relative to primiparous rats (p = 0.023; **Fig. 8A**). There was also a significant main effect of time (p<0.001), but not reproductive status (p=0.14) nor an interaction (p=0.325).

**Figure 8.**
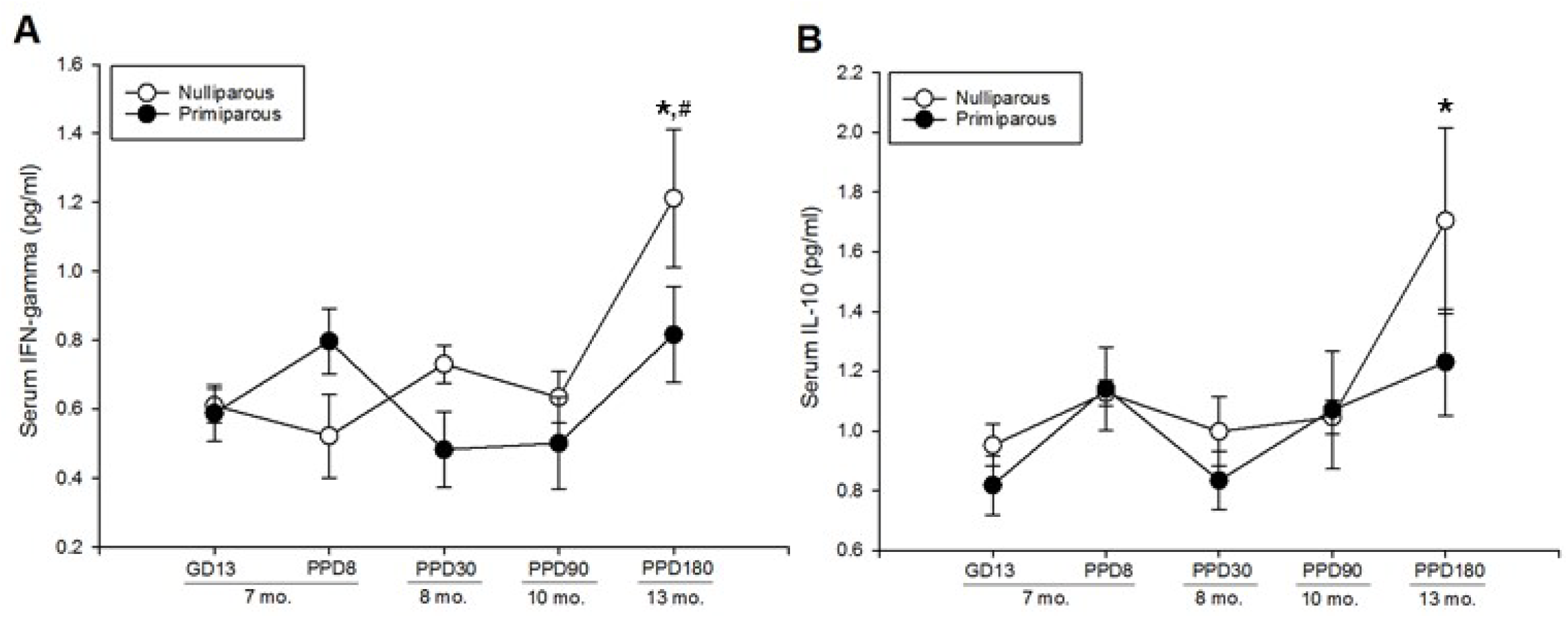
Serum levels of interferon-**γ (A)** and interleukin-10 **(B)** in primiparous and nulliparous rats. The x-axis represents time relative to gestation and parturition in primiparous groups, and approximate age in months. There was a significant ageing-related increase in IFN-γ **(A)** and IL-10 **(B)** in nulliparous but not primiparous rats. **\*** indicates p <0.004, significantly different from all other nulliparous groups. # indicates p = 0.023, significantly different from13-month-old primiparous rats. Data are represented in mean values ± SEM. IFN-γ = interferon gamma, IL-10= interleukin 10, GD= gestation day, PPD= postpartum day.

Similarly, serum IL-10 increased significantly with age in nulliparous but not primiparous rats; significantly higher levels of IL-10 were detected in 13-month old nulliparous rats relative to all other nulliparous groups (P<0.002; **Fig. 8B**), but no significant differences were found between any of the primiparous groups (all p’s>0.05; planned comparisons; **Fig. 8B**). There was also a significant main effect of time (p<0.003), but not reproductive status (p=0.11) nor an interaction (p=0.45).

### 3.10. Serum IL-4 was transiently increased in the early postpartum then persistently reduced by parity

Nulliparous rats had higher levels of IL-4 than primiparous rats, regardless of time point (main effect of reproductive status: F(1, 38)=7.63, p<0.009, **Fig. 9A**). Regardless of reproductive status, there was an age-related increase in serum IL-4, with significantly elevated levels at 13 months compared to all groups at 8 months of age and younger (all p’s< 0.04; main effect of time: F(4, 38)=4.29, p<0.006). There was no significant reproductive status by time interaction (F(4, 38)=1.73, p=0.16), but a priori we expected cytokine levels to be altered in the early postpartum period in primiparous rats. Indeed, primiparous rats showed a trend for a transient increase in serum IL-4 in the early postpartum period, with higher levels at PPD8 relative to GD13 (p=0.029) and PPD30 (p=0.027; a priori comparisons, missing significance with correction; **Fig. 9A**). In age-matched nulliparous control groups, serum IL-4 was not significantly different in 7.5-relative to 7-or 8-month-old rats (p’s >0.45).

**Figure 9.**
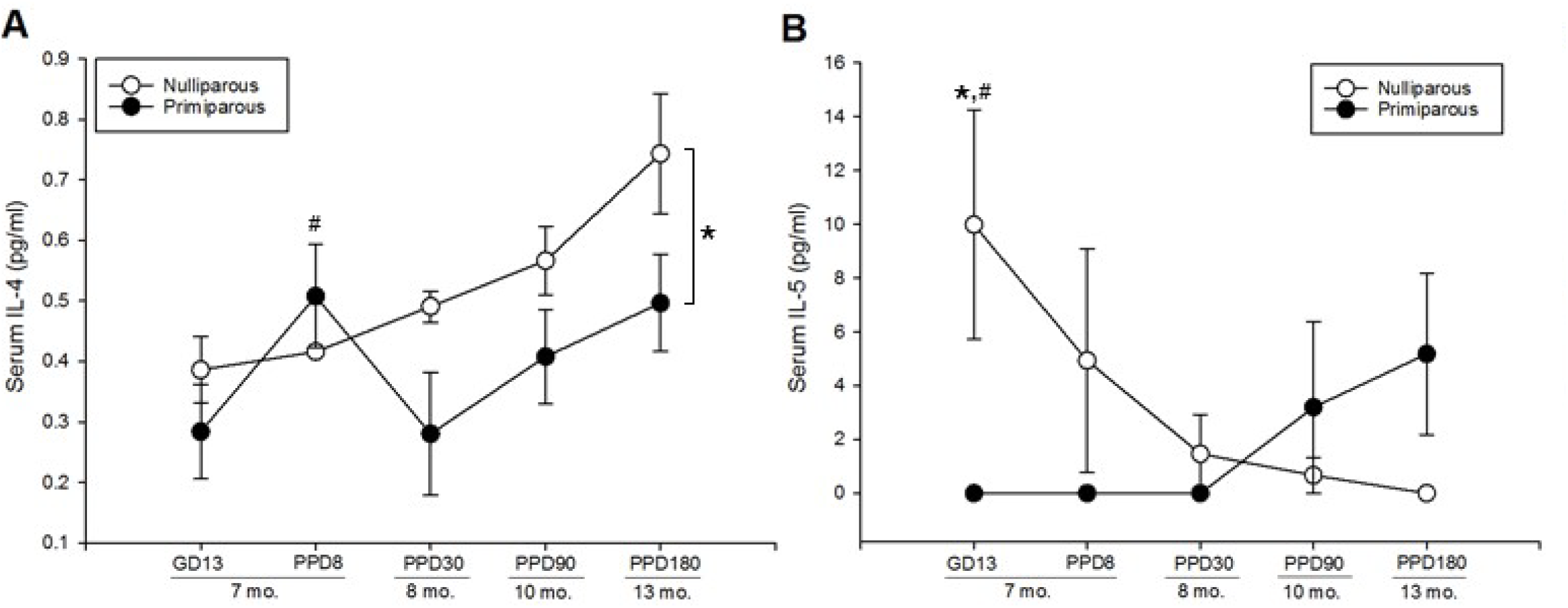
Serum levels of interleukin-4 **(A)**, and interleukin-5 **(B)** in primiparous and nulliparous rats. The x-axis represents time relative to gestation and parturition in primiparous groups, and approximate age in months. **(A)** Serum IL-4 levels were transiently increased in primiparous rats at PPD8, but persistently suppressed by parity thereafter. # indicates p<0.03, trend towards significance relative to GD13 and PPD30. * indicates p<0.009, main effect of primiparity to reduce IL-4 levels **(B)** Serum IL-5 was transiently blunted during gestation, and a significant age-related decline in serum IL-5 was found only in nulliparous rats. * indicates p=0.006, significantly higher than primigravid rats at GD13. # indicates p<0.02, significantly higher than all nulliparous groups between 8 and 13 months of age.

### 3.11. Serum IL-5 was transiently reduced during gestation and showed an age-related decline in nulliparous but not primiparous rats

Serum IL-5 levels were significantly reduced in gestation (GD13), relative to age matched nulliparous controls (p=0.006; time by reproductive status interaction, F(4, 37)=3.31., p=0.020; **Fig. 9B**). Further, IL-5 levels declined with age in nulliparous animals, as higher levels were detected at 7 months relative to 8, 10 and 13 months (all p’s<0.02; **Fig. 9B)**. Although non-significant, IL-5 levels increased in primiparous animals with age, suggesting a reversed pattern of age-related changes in IL-5 compared to nulliparous rats. There were no significant main effects of time or reproductive status (all p’s >0.1).

### 3.12. Serum IL-13, IL6, CXCL1, IL-1β, and TNF-α, were not significantly altered by parity or age

Regardless of reproductive status, there was a trend towards significance for a main effect of time to affect IL-13 (F(4, 37)=2.46, p=0.06; **Fig. 10A**), where IL-13 levels were elevated at 13 months compared to 8 months (p <0.002; planned comparisons). There were no significant main effects of reproductive status or 17β-estradiol (covariate), and no time by reproductive status interaction (all p’s >0.4). There were no significant main effects of reproductive status or age, nor an interaction for serum concentrations of IL-6 (all p’s > 0.21, **Fig. 10B**), CXCL1 (all p’s >0.40 **Fig. 10C**), or IL-1β (all p’s > 0.32, **Fig. 10D**). There were trends towards significance for a main effect of parity to increase TNF-α (p = 0.064) and a main effect of time (p = 0.061) for TNF-α to decline with age, but no significant reproductive status by time interaction (p = 0.72. **Fig. 10E**).

**Figure 10.**
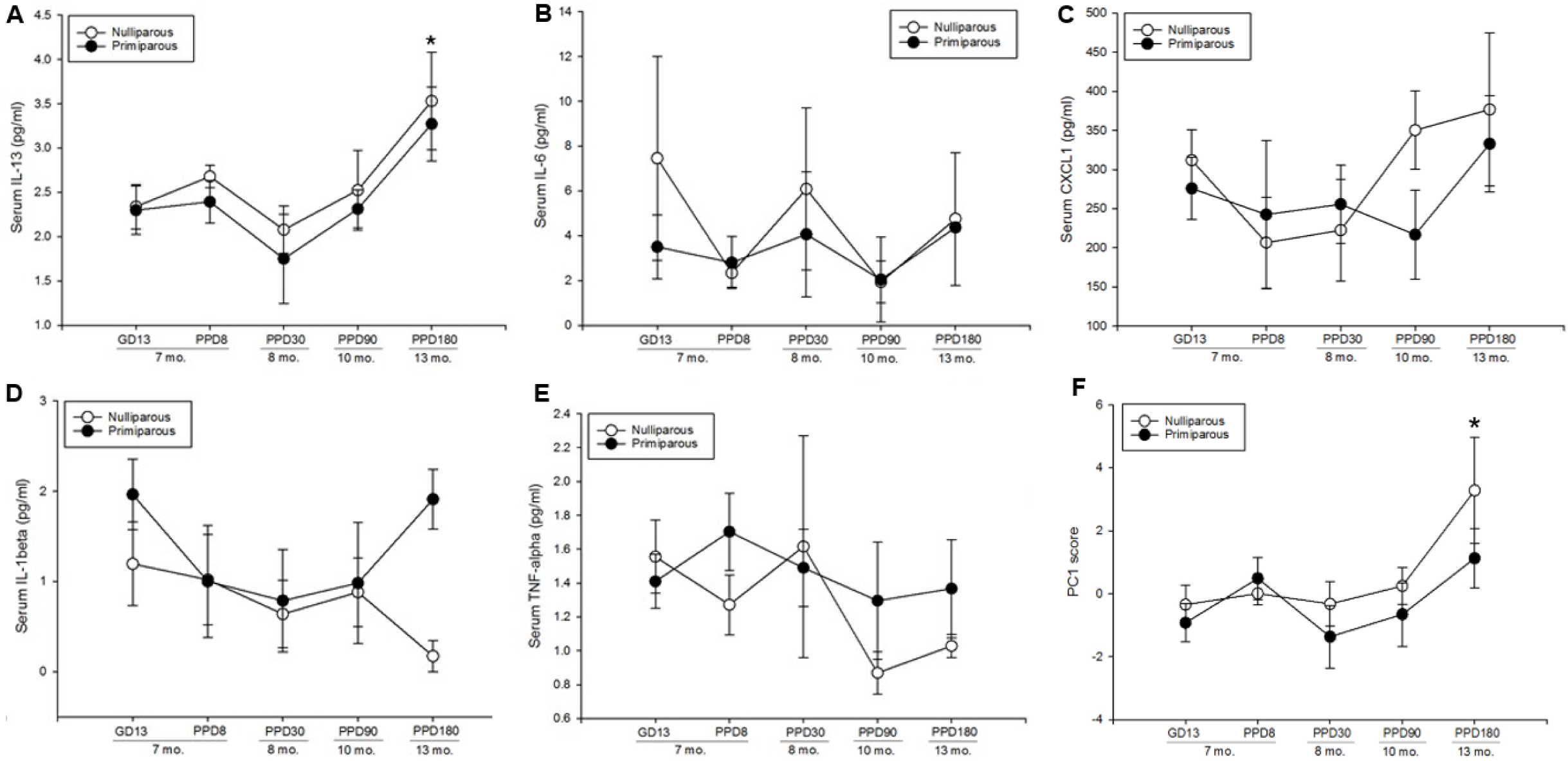
Serum concentrations of interleukin-13 **(A)**, interleukin-6 **(B),** CXCL1 **(C)**, interleukin-β **(D)**, and tumor necrosis factor alpha-α **(E)** in primiparous and nulliparous rats. The x-axis represents time relative to gestation and parturition in primiparous rats, and approximate age in months. **(A)** There was a trend towards significance for serum IL-13 to increase with age regardless of reproductive status. * indicates p<0.002 significantly different from 8-month old rats. Reproductive status and age had no significant effects on serum levels of interleukin-6 **(B),** CXCL1 **(C)**, and interleukin-β **(D)**. **(E)** There were trends towards significance for parity to increase (p= 0.064) and for age to decrease (p= 0.061), TNF-α concentrations. **(F)** Principal Component 1 scores in primiparous and nulliparous rats, * indicates p’s<0.007, significantly higher PC1 scores in 13-relative to 7- and 8-month-old nulliparous rats. Data are represented in mean values ± SEM. GD= gestation day, PPD= postpartum day. Data are represented in mean values ± SEM. IL-4= interleukin-4; IL-5 = interleukin-5; IL-13 = interleukin-13; GD= gestation day; PPD = postpartum day; mo.= age in months.

### 3.13. Principal Component Analysis of serum cytokines

The model generated 4 principal components, which accounted for 82.6% of the variance within the dataset, with the first principal component explaining 40.4% of the variance, the second 18.2%, the third 12.6%, and the fourth 11.4%. Interestingly, IFN-γ, IL-10, IL-13, and IL-4 loaded heavily onto Principal Component 1 (PC1; see **Table 3**). These same cytokines also showed the most robust alterations with parity and age with ANOVA, therefore the PCA ultimately verified our individual ANOVA analyses. Subsequently, we analyzed PC1 scores using ANOVA, which revealed a main effect of time (F(4, 37)=3.78, p=0.011; **Fig. 10F**), with higher scores at 13 months of age relative to all other age groups (all p’s <0.033). Planned comparisons reveal a more robust age-related increase in PC1 scores in nulliparous rats, with higher scores in 13-relative to 7- and 8-month-old rats (p’s<0.007). In contrast, there were no significant differences in PC1 scores between any primiparous groups (p’s>0.038; non-significant due to Bonferroni correction). There was no significant main effect of reproductive status, nor an interaction (p’s >0.11). Interestingly, the pattern observed here is akin to the age-related increase in IL-10, IFN-γ, and IL-4 in nulliparous rats, obtained with individual ANOVA analyses. Further, IL-6 and TNF-α loaded heavily onto PC2, and IL-1β and IL-5 loaded heavily onto PC3 and PC4, respectively (Table 1).

**Table 3.**
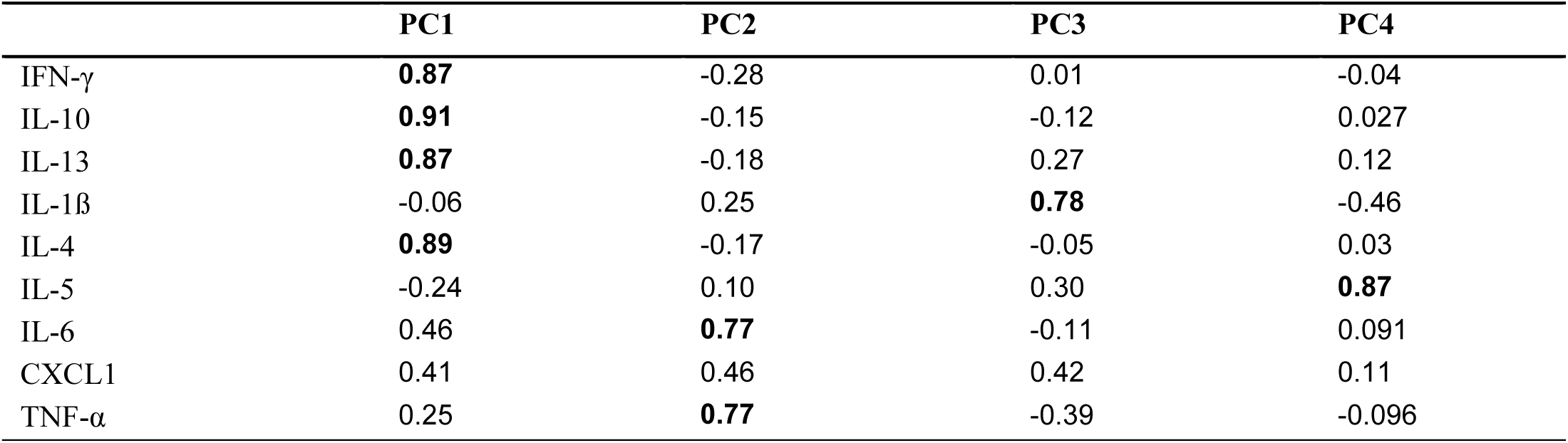
Principal Component Analysis loading table.

### 3.14. Reproductive status influenced the direction and magnitude of correlations between microglia and neurogenesis measures

Previous studies indicate that microglia play a role in adult hippocampal neurogenesis under baseline and inflammatory conditions (Ekdahl et al., 2003; Monje, 2003; Sierra et al., 2010). However, the role of microglia in motherhood-associated neurogenic changes have not been studied, thus, to indirectly examine this we correlated our measures of neurogenesis and microglia. Increased average length of processes was significantly correlated with a higher number of Ki67-IR cells in nulliparous (r = 0.58, p = 0.005; **Fig. 11A**), but not primiparous rats (r = 0.089, p = 0.67), and these correlations were significantly different (z = 1.81, p = 0.035). Similarly, increased average length of processes was significantly correlated with a higher number of DCX-IR cells in nulliparous (r = 0.62, p = 0.002; **Fig. 11B**), but not primiparous rats (r = 0.11, p = 0.60; **Fig. 11B**), and these correlations were significantly different (z = 1.96, p = 0.025). Interestingly, increased Ki-67-IR cell number was significantly associated with more DCX-IR cells in nulliparous rats (r = 0.86, p<0.001; **Fig. 11C**), but not in primiparous rats (r = 0.30, p = 0.15; **Fig. 11C**), and the difference between the correlations was significant (z = 3.04, p = 0.001).

**Figure 11.**
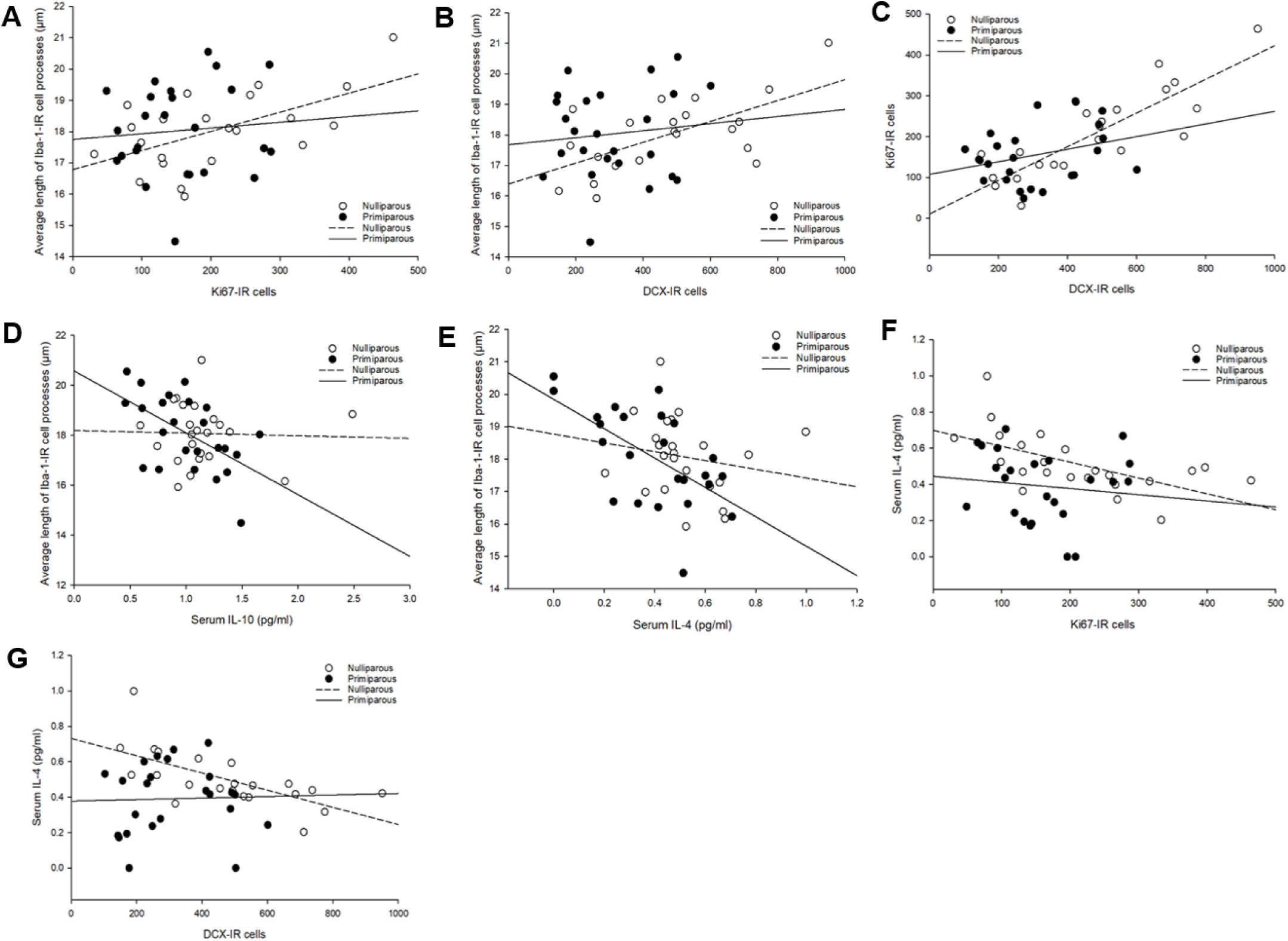
Correlations between dependent variables of interest. Increased average length of Iba-1-immunoreactive (IR) cell processes was associated with higher Ki67 **(A)** and doublecortin (DCX; **B**) expression in nulliparous rats only. KI67-IR cell number was significantly and positively correlated with DCX-IR cell number in nulliparous rats only **(C)**. Elevated serum concentrations of IL-10 **(D)**, and IL-4 **(E)** were significantly associated with shorter average Iba-1-IR cell processes in primiparous rats only. In nulliparous rats only, increased Ki67-IR **(F)** and DCX-IR **(G)** cell number was significantly associated with lower serum IL-4.

### 3.15. Reproductive status influenced the direction and magnitude of the correlations between circulating cytokine concentrations and neural measures

Peripheral cytokine signals propagate to the brain and can influence neuroimmune function (Miller et al., 2014; Quan and Banks, 2007), thus we examined correlations between serum cytokine levels and microglial measures to assess whether central and peripheral inflammatory indicators are associated in parous and non-parous rats. Interestingly, in primiparous rats, the average length of Iba-1-IR cell processes was negatively correlated with serum IL-10 (r = −0.56, p = 0.005; **Fig. 11D**), and IL-4 (r = −0.6, p = 0.002; **Fig. 11E**). On the other hand, length of cell processes was not significantly correlated with IL-10, or IL-4 in nulliparous rats (all p’s > 0.36). These correlations were significantly different between reproductive status groups for IL-10 (z = 2.13, p = 0.017), but not for IL-4 (p = 0.054). Regardless of reproductive status, no other significant correlations were found between length of Iba-1-IR cell processes and all other measured cytokines (all p’s > 0.056; all trends towards significance appear in primiparous groups only). Further, there were no significant correlations between any of the cytokines and Iba-1-IR cell density, regardless of reproductive status (all p’s > 0.23).

In nulliparous rats, increased serum concentrations of IL-4 were associated with fewer DCX-IR and Ki67-IR cells (Ki67: −0.57, p=0.006; **Fig. 11F**; and DCX: r= −0.70, p<0.001; **Fig. 11G**). There were no significant correlations between any cytokines and Ki67- or DCX-IR cells in primiparous rats (all p’s > 0.19). These correlations were significantly different between reproductive status groups in both instances (DCX: z = −2.58, p < 0.005; Ki67: z = −1.82, p = 0.034).

### 3.16. Serum 17β-estradiol concentrations were not significantly correlated with hippocampal cell proliferation, Iba-1 measures, or serum cytokine concentrations

Serum 17β-estradiol concentration were not significantly associated with Ki67 expression in primiparous or nulliparous rats (p’s > 0.1). Regardless of reproductive status, there were also no significant correlations between 17β-estradiol concentrations and any measure of Iba-1-IR cells (density, cell body size, average length and number of processes; all p’s >0.1), or any of the cytokine concentrations (all p’s >0.17).

## 4. Discussion

Here, we report short- and long-term effects of maternal experience on hippocampal neurogenesis, microglial density and morphology in the dentate gyrus, and circulating cytokine levels, culminating six months after parturition. We found that adult hippocampal neurogenesis was suppressed in mid-gestation and up to one month postpartum. Interestingly, the ageing trajectory of neurogenesis was modulated by reproductive experience, as neurogenesis levels declined from 7 to 13 months of age in nulliparous rats, but showed a slight increase in primiparous rats across the same period. Hippocampal cell proliferation was suppressed in mid-gestation and the early postpartum period in primiparous rats, but normalized thereafter, as an age-related decline in cell proliferation was observed regardless of previous parity. Further, microglia in the dentate gyrus displayed a more activated morphology in the early postpartum period, followed by a transient increase in microglial density in the later postpartum period and overall smaller microglia soma size in primiparous compared to nulliparous rats. We found alterations in circulating cytokine levels during pregnancy and the early postpartum period, and more intriguingly, we show that the age-related changes in circulating cytokine levels were dependent on parity. We also observed that reproductive status shifted the associations between microglia and neurogenesis, with average length of Iba-1-IR cell processes being positively associated with neurogenesis in nulliparous but not primiparous. Further, parity modulated the correlations between serum cytokines and microglial morphology, and between serum cytokines and neurogenesis levels. These correlations suggest that the relationships between immune processes and neurogenesis may be modified with parity. Lastly, 17β-estradiol concentrations were reduced in the early postpartum period (PPD8) and in mid-gestation (GD13), in line with previous findings (Rosenblatt et al., 1988). Beyond the expected pregnancy and postpartum changes in circulating 17β-estradiol, we found that concentrations were not significant moderators of most variables, including hippocampal cell proliferation, microglial density and morphology, and cytokine concentrations. Collectively our data suggest that maternal experience has transient and delayed effects on hippocampal neurogenesis, microglia, and the peripheral inflammatory milieu.

### 4.1. Adult hippocampal neurogenesis was suppressed during gestation and the postpartum period

We report that adult hippocampal neurogenesis, measured via the expression of doublecortin, was suppressed beginning in mid-gestation in primigravid rats. Few studies to date have examined the survival of new cells in the maternal hippocampus during gestation (Pawluski et al., 2010; Rolls et al., 2008). Our findings are, however, consistent with a study that found suppressed neurogenesis in pregnant mice during mid- and late-gestation (Rolls et al., 2008). In contrast, the expression of PSA-NCAM was increased in the dentate gyrus of pregnant rats at GD18 (Banasr et al., 2001), indicating a potential increase in neurogenesis, as PSA-NCAM is expressed on newly generated and migrating neurons (Rutishauser, 2008). However, because PSA-NCAM is also expressed on neurons undergoing other forms of plasticity (Rutishauser, 2008), its expression provides limited and non-specific information regarding neurogenesis levels. Another study in rats found that the survival of cells produced on gestation day 1 was not significantly altered when examined across gestation (Pawluski et al., 2010), partially contrasting with our current findings. Importantly, however, DCX is expressed in immature neurons between 2 hours and 21 days after production (Brown et al., 2003). Therefore, our current data provide information on the population of cells produced as early as 8 days prior to impregnation, and as late as the day of euthanasia (GD13). Therefore, inconsistencies between the two studies are not surprising, considering that the cell populations examined were produced under different conditions. We also observe a concurrent reduction in cell proliferation on GD13 indicating that this may underlie the decline in immature neurons at this time. Although no prior studies have examined cell proliferation in mid-gestation, cell proliferation was not altered on GD1 (Pawluski et al., 2010), GD7 (Shingo et al., 2003), or GD21 (Furuta and Bridges, 2005). Therefore, a more detailed time-course analysis of hippocampal plasticity during pregnancy is warranted.

Consistent with a prior study (Workman et al., 2015), we also found reduced DCX expression in the postpartum period, evident until PPD30. This finding is also in keeping with past work showing reduced survival in new cells labelled on PPD2 and examined 21 days later (Pawluski and Galea, 2007), and in new cells labelled in mid-gestation (GD11-12) and examined 14 days later, in the early postpartum period (Rolls et al., 2008). We found reductions in hippocampal cell proliferation in the early postpartum period, which normalized by PPD30, in line with previous data (Darnaudéry et al., 2007; Leuner et al., 2007; Pawluski and Galea, 2007; Rolls et al., 2008). Thus, the suppression in immature neurons found at PPD8 likely resulted from a reduction in both cell proliferation and survival, whereas the suppression at PPD30 is likely due to decreased cell survival rather than proliferation. Although the functional significance is not known, suppressed neurogenesis in the maternal brain may be mechanistically associated with the enhanced susceptibility to mood disorders in the peripartum period (Hendrick et al., 1998). Further, separate lines of evidence indicate that adult neurogenesis is involved in hippocampal regulation of the HPA axis at least in males (Schloesser et al., 2009; Snyder et al., 2011), and that the HPA axis undergoes substantial adaptations during pregnancy and the postpartum (De Weerth and Buitelaar, 2005; Lightman et al., 2001; Slattery and Neumann, 2008). Therefore, reductions in neurogenesis in the maternal hippocampus could influence HPA axis function. Suppressed neurogenesis may also be linked to deficits in hippocampus-dependent learning and memory reported in late pregnancy and the early postpartum period (reviewed in (Workman et al., 2012)).

### 4.2. Maternal experience altered the trajectory of age-related changes in hippocampal neurogenesis

Between 8 and 13 months of age, immature neurons in the dentate gyrus declined significantly in nulliparous rats but showed a slight increase in primiparous rats. We observe an age-related decline in cell proliferation (Ki67-IR cells) regardless of reproductive status, suggesting that the differential effects in immature neurons (DCX-IR cells) are driven by differences in cell survival. Previous work indicates that hippocampal neurogenesis steadily declines with age, with the most substantial decline occurring between adulthood and middle age in female rats (Driscoll et al., 2006; Kuhn et al., 1996; Nacher et al., 2003), consistent with our current data from nulliparous rats. Thus, the increase in neurogenesis levels in middle-aged primiparous rats suggests that reproductive experience can modify the trajectory of age-related alterations in neurogenesis. Alternatively, it may also be reasonable to interpret this finding as merely a normalization of neurogenesis to nulliparous levels. However, two previous reports indicate higher neurogenesis levels in primiparous and multiparous relative to nulliparous middle-aged rats (Barha et al., 2015; Galea et al., 2018), and as such an altered aging trajectory is conceivable. It is possible that a more robust difference in neurogenesis levels would arise only after multiple reproductive experiences or later into middle age. There is emerging evidence from human and rodent studies suggesting that motherhood can alter the course of age-related cognitive decline (Beeri et al., 2009; Colucci et al., 2006; Cui et al., 2014; Gatewood et al., 2005). For example, reproductive experience mitigated the age-related decline in spatial memory in rats (Gatewood et al., 2005) and mice (Cui et al., 2014). Other studies indicate that parity is associated with cognitive impairment in the ageing female (Beeri et al., 2009; Colucci et al., 2006). These inconsistencies may be reconciled by more complex interactions with genetic factors that have been associated with pathological cognitive ageing (Corbo et al., 2007; Cui et al., 2014). While speculative, the modest increase in hippocampal neurogenesis in middle-aged primiparous rats may be associated with enhanced hippocampus-dependent cognition that is seen at that time.

Interestingly, we observed differences in the relationship between levels of cell proliferation and immature neurons depending on reproductive status, such that a significant positive correlation between the two measures was only seen in nulliparous rats. In addition, increased IL-4 concentrations were associated with reduced proliferation and immature neurons in the hippocampus of nulliparous but not primiparous rats. Previous work points to a role of IL-4 in the regulation of cell proliferation under conditions of neurodegeneration (Bhattarai et al., 2016), therefore our findings suggest that this role may be altered by parity. These relationships should be further investigated, as they appear when reproductive status groups are collapsed across age, but nonetheless indicate that parity may modulate the effects of immune signaling on hippocampal neurogenesis.

### 4.3. Microglia assumed a de-ramified morphology in the early postpartum period

Microglia display a predominantly ramified morphology under basal conditions, and de-ramification is thought to be indicative of increased classical activation under inflammatory conditions (Luo and Chen, 2012). Here, we show that microglia in the dentate gyrus exhibited significantly shortened processes at 8 days postpartum, suggesting an increase in microglial activation in the early postpartum period. This de-ramification was a transient morphological modification as the average length of processes was not significantly different from nulliparous controls by PPD30. We further show a small but significant shift in the percentages of microglia assuming different morphological states at PPD8. Specifically, we observe a reduction in ramified morphology and an increase in stout morphology, which in tandem with our data on length of processes indicates a shift towards classical microglial activation in the dentate gyrus in the early postpartum period. However, we interpret these findings with caution, as the information that can be deduced from morphological phenotype about functional states is limited (Boche et al., 2013). 17β-estradiol concentrations significantly moderated the average length of microglial processes, suggesting that the observed de-ramification at PPD8 might be moderated by postpartum reductions in estradiol. However, as this measure of estradiol concentrations provides information about a small window of time around perfusion, a complete picture of the role of estradiol cannot be extrapolated from the current study. Future studies should also consider the role of other hormones, including progesterone and corticosterone, in motherhood-associated changes microglia.

To our knowledge, only two studies to date have examined microglia in the maternal brain (Haim et al., 2017; Posillico and Schwarz, 2016), with findings partially consistent with our current data. Haim et al. (2017) reported a decrease in the number of microglia with a ramified morphology on postpartum day 8, in several regions including the dorsal hippocampus. This is in line with our current report of microglial de-ramification and a shift in percentages of morphological states in the dentate gyrus on the same postpartum day (PPD8). In the current study, we found an increase in the density of microglia in the dorsal and ventral dentate gyrus at PPD30, but no alteration in density during gestation or the early postpartum period. While no other studies have examined microglia in the maternal brain as late as 30 days postpartum, our finding that microglial density in the hippocampus remains unchanged during gestation and the early postpartum period contrasts previous reports (Haim et al., 2017; Posillico and Schwarz, 2016). These previous studies found reduced microglial density in several brain regions from late gestation to the early postpartum (GD20, and PPD1, 8, and 21: Haim et al., 2017; PPD0: Posillico and Schwarz, 2016). Further, Haim and colleagues (2017) found that microglial density normalized to nulliparous control levels by PPD21 in all regions examined except the dorsal hippocampus. It appears, however, that the inconsistencies in findings may be accounted for by differences in microglial densities within sub-regions of the dentate gyrus, or by methodological differences related to density measurement. For example, Haim et al. (2017) examined Iba-1 density within the dorsal dentate gyrus only, whereas we included samples from both the dorsal and ventral dentate gyrus. In addition, as we were primarily interested in the neurogenic niche, we quantified Iba-1-IR cells within the GCL, the SGZ, and a thin band of the ML, whereas Haim and colleagues did not specify sub-regions within the dentate gyrus. Finally, Haim et al (2017) utilized optical density, whereas density here was defined as the number of cells per volume of dentate gyrus. Interestingly, the increase in microglial density that we find at PPD30 coincides with a return to normalized microglial morphology. We speculate that this may represent a resolution from the pro-inflammatory state at PPD8. Overall, our novel data provide an important addition to the literature indicating that pregnancy-related immune adaptations are not limited to the periphery, but also exist in the brain. More specifically, our data indicate the existence of a pro-inflammatory hippocampal environment in the early postpartum period. Importantly, increased microglial activation may be central to the pathophysiology of depression (Kreisel et al., 2014; Miller and Raison, 2015; Setiawan et al., 2015), thus it is conceivable that similar processes are implicated in postpartum depression. Our current findings which point to an increase in microglial activation in the early postpartum may represent a neural mechanism of enhanced susceptibility to mood disorders at that time, however additional work would be required to directly test this.

The effects of parity on microglial morphology were not limited to the early postpartum period. Specifically, although the number of microglial cell processes increased significantly with age in nulliparous rats, this effect was prevented by parity. Further, parity reduced microglial soma size, an effect which appears to emerge at PPD30 onwards. The functional significance of these alterations cannot be determined from the current study, but as increased microglial soma size is indicative of classical activation, it is possible that parity may dampen microglial activation in the ageing brain. To gain better insight into the functional significance of these changes, future studies should investigate how previous parity may impact microglial structure and function in the ageing brain in response to an immune challenge.

Interestingly, we found that reproductive status affected the associations between microglia and neurogenesis in the hippocampus. Specifically, increased cell proliferation and immature neurons were significantly associated with more ramified microglial morphology in nulliparous rats only. On the other hand, increased cell proliferation and immature neurons were associated with higher microglial density in primiparous rats only. These observations suggest that the neuroimmune regulation of adult hippocampal neurogenesis is affected by reproductive status. Importantly, in addition phagocytic activity during development and disease, microglia are important players in the regulation of adult hippocampal neurogenesis, where they phagocytose apoptotic new cells, while maintaining ramified morphology and a non-inflammatory environment (Sierra et al., 2010). Thus, future research should directly examine the possibilities of microglial phagocytosis in the regulation of neurogenesis during pregnancy and the postpartum period.

### 4.5. Maternal experience modifies the age-related changes in circulating cytokine levels

In addition to expected cytokine alterations during pregnancy and the early postpartum period (Holtan et al., 2015; Shimaoka et al., 2000), we observed both persistent and delayed effects of reproductive experience on the circulating cytokine profile. Specifically, after an initial increase at PPD8, IL-4 levels were persistently blunted in primiparous rats relative to nulliparous controls. Further, in nulliparous but not primiparous rats, IFN-γ and IL-10 increased and IL-5 declined significantly with age. Adaptations to the immune systems during pregnancy and the early postpartum period are well established (PrabhuDas et al., 2015). On the other hand, little attention has been paid to potential long-term effects of motherhood on the immune system. To our knowledge, only a few studies have examined the effect of parity on immune systems in aged female mice and one in aged rats. Together these studies suggested that parity may delay certain indicators of immune senescence, as they relate to alterations in cytokine production *in vitro* from activated spleen cells (Barrat et al., 1997a), and the distribution of immune cell populations in the spleen (Barrat et al., 1997b), and bone marrow (Barrat et al., 1999). Another study from our laboratory found a trend for increased serum levels of IL-6 in primiparous compared to nulliparous rats at 15 months of age (Galea et al., 2018). Here, we report that a single reproductive experience alters serum cytokine levels when examined in middle age, up to six months after the reproductive event.

Importantly, peripheral cytokines can affect brain function, as they can access the central nervous system through various mechanisms, including active and passive transport, and the activation of cytokine receptors on afferent nerve fibers (reviewed in Miller et al., 2014; Quan and Banks, 2007). Thus, the effects of parity to modify age-related changes in peripheral cytokines may have ramifications for the ageing brain in general, and more specifically the hippocampus, as it contains one of the highest densities of proinflammatory cytokine receptors in the brain (reviewed in Loftis et al., 2010). Inflammation is a core characteristic of the ageing processes, and the pro-inflammatory cytokine IFN-γ increases with age (Oxenkrug, 2011; Rodríguez et al., 2007). Thus, our current data suggest that parity may prevent or delay at least certain aspects of ageing-related inflammation. Interestingly, from PPD30 onwards, we observed a sustained suppression in serum IL-4 in primiparous rats relative to nulliparous controls. Traditionally considered an anti-inflammatory cytokine (Hart et al., 1989), elevated IL-4 in nulliparous rats may be suggestive of a compensatory response to attenuate a pro-inflammatory state that is indicated by elevated IFN-γ. However, IL-4 is pleiotropic (Milner et al., 2010), and as such also can have pro-inflammatory properties. For example, sustained exposure to elevated levels of IL-4 was associated with increased inflammation (Milner et al., 2010). The same study found prolonged IL-4 exposure to be associated specifically with elevated levels of IL-10 and IFN-γ, but not IL-6 and TNF-α. This is in line with our current data in which IL-4, IL-10, and IFN-γ were concurrently elevated in middle-aged nulliparous rats. Thus, the cytokine profile in nulliparous groups may be alternatively driven by an increase in IL-4. We investigated correlations between peripheral cytokines and microglial morphology as a plethora of work suggests that systemic inflammation can profoundly affect microglial activation (Reviewed in Hoogland et al., 2015). Interestingly, we found that higher concentrations of IL-10 and IL-4 were associated with more de-ramified microglial morphology in primiparous but not nulliparous rats. The mechanisms and consequences of these altered relationships between peripheral cytokines and microglial morphology are not known, and these findings should be interpreted with caution, as the correlations were detected when age was not included as a factor. In the future, it is also important to examine whether parity may have similar effects on age-related changes in brain cytokines, particularly in the hippocampus. While the functional consequences of the observed differences in cytokine profiles cannot be determined from our current finding, we demonstrate here that the trajectory of immune senescence is altered by parity.

## 5. Conclusions

In summary, we report that maternal experience suppressed hippocampal neurogenesis (proliferation and immature neurons) during gestation and the postpartum period and mitigated the decline in neurogenesis in middle age (immature neurons). Maternal experience also resulted in transient microglial de-ramification in the dentate gyrus, suggesting the existence of a pro-inflammatory hippocampal environment in the early postpartum period. In addition to short-term cytokine alterations, maternal experience modified the trajectory of age-related changes in circulating cytokine levels. These findings should encourage future work aimed at delineating the functional consequences for behaviour and immune function across the peripartum period and beyond, especially in relation to maternal mood and cognition. Importantly, our data provide support for the notion that female reproductive history should be regarded as an important determinant of ageing-related changes in physiology.

## Acknowledgments

The Authors thank Dr. Timothy Kieffer and Travis Webber for generously providing access to their Sector Imager, and Arianne Albert for her assistance with principal component analyses.

